# Microbiota during early life windows orchestrates intestinal maturation and barrier formation in the offspring

**DOI:** 10.64898/2026.09.24.754204

**Authors:** Cristina Kalbermatter, Sandro Christensen, Irene Keller, Nerea Fernandez Trigo, Joana Carlevaro-Fita, Dorothée L. Berthold, Stephanie Talker, Anna Wenning, Rémy Bruggmann, Uwe Sauer, Mercedes Gomez de Agüero, Andrew J. Macpherson, Stephanie C. Ganal-Vonarburg

## Abstract

There is a critical window in early life for immune development that prepares a young mammal for postnatal exposure to environmental, dietary and microbial antigens. Both the maternal microbiota and endogenous colonization have emerged as pivotal drivers in this process. Using gnotobiotic mouse models and reversible colonization to model the effects of the maternal microbiota during pregnancy compared with early life colonization on the small intestinal epithelium we show that both microbial metabolites originating from the maternal microbiota and the postnatal endogenous microbiota drive intestinal epithelial maturation by modulating epigenetic signatures. Microbiota-derived signals from both the pregnant mother and from endogenous postnatal colonisation drove the BLIMP1-guided epithelial neonatal-to-adult transition at weaning through epigenetic mechanisms and direct transcriptional effects. These microbial signals also shaped the developmental trajectory of barrier-forming tight junctional claudins in the neonatal intestine, increased aerobic respiration, and increased the gene expression of xenosensors. Thus, both maternal and endogenous microbiota derived drivers of the offspring’s epigenetic and transcriptomic intestinal epithelial landscape fine-tuned the timing of epithelial maturation and augmented functional plasticity at weaning.

## INTRODUCTION

The earliest stages of life play a pivotal role in shaping the intestinal microbiota and subsequent immune system development, designated as “windows of opportunity” ^1^. Perturbations in the microbiota during early life can increase susceptibility to allergies, obesity and autoimmunity during adulthood ^2^. Successive postnatal colonization with mutualistic microbes is first seeded by bacteria in the birth canal. Although the fetus emerges from a protected and sterile uterine environment ^3^, it has already been exposed to transplacental metabolites originating from the maternal microbiota and diet ^4–7^. Using reversible microbial colonization exclusively during pregnancy we showed that germ-free pups, that had only ever been indirectly exposed to metabolites of the gestational maternal microbiota, had enhanced numbers of innate immune cells in the intestinal lamina propria, and an altered small intestinal transcriptome compared to offspring born to non-exposed dams ^4^. This suggested that a window of epigenetic or regulatory opportunity may even originate independently of endogenous colonization.

The microbiota is recognized to drive differential gene expression of intestinal epithelial cells ^8–10^. The intestinal epithelial layer comprising differentiating stem cells, enterocytes, goblet cells, Paneth cells, enteroendocrine cells and Tuft cells is only indirectly exposed via the placenta to the molecular diaspora from the commensal microbiota and dietary components before birth but becomes directly and progressively exposed following postnatal lactation.

During early postnatal development, the neonatal epithelium is characterized by vacuolated enterocytes and a scarcity of mature goblet cells and Paneth cells, resulting in a thinner mucous layer and scarce antimicrobial peptides compared to the adult epithelium ^11–13^. Whereas pore-forming claudins are characteristic of the murine neonatal epithelium ^14–16^, as the infant approaches weaning, vacuolated enterocytes diminish, the brush border surface increases, and barrier-forming claudins appear, transitioning towards an adult-like epithelium ^11,17–19^. During this transition, the epithelium enhances its metabolic capacity and adjusts its sensory and antimicrobial strategies by increasing antimicrobial peptide production and modulating Toll-like receptor (TLR) expression ^20–22^. Since the diet, the microbiota and epithelial adaptation all change from birth to weaning, the timing and mechanisms of the microbiota interactions before and after birth are crucial to understand microbial influences on epithelial development.

Direct effects on epithelial transcription by the microbiota or other environmental and dietary molecules take place in the epigenetic landscape at each stage of development and differentiation. The maturation of the epigenetic landscape, such as alterations in DNA cytosine methylation and covalent histone modifications, may be not only canonically programmed according to age, but also driven by this molecular exposure that induces durable changes in gene expression within intestinal epithelial cells during early life. DNA methylation gains in intestinal stem cells from birth to weaning have been found to be aligned with barrier integrity ^23,24^. That the intestinal epithelial chromatin architecture can be shaped by the microbiota as well as developmentally programmed has been shown by comparing DNA methylation landscapes and gene expression in small intestinal epithelial cells of germ-free and colonized mice at different stages of life ^23,25,26^ and may be determined very early after birth. However, the specific role of the maternal microbiota uncoupled from postnatal environmental exposures and endogenous colonization, or how these influences integrate, has remained unclear.

In this study we show that both maternal and endogenous intestinal microbial colonization during these early life windows drive intestinal epithelial cell development. We demonstrate the effect of maternal intestinal microbes on gene expression and epigenetic landscapes shaping maturation, barrier formation and metabolic adaptation of the postnatal intestinal epithelium.

## RESULTS

### Maternal and postnatal colonization shape the epigenetic and transcriptional program of small intestinal epithelial cells early in life

We previously reported on maternal colonization during pregnancy shaping intestinal gene expression in the offspring at postnatal day 14 ^4^. A large proportion of altered genes were of epithelial origin, yet the underlying mechanisms and durable consequences within the epithelial compartment remained poorly understood. We postulated that microbe-host interactions that are initiated in the pregnant maternal gut may have a lasting influence on the postnatal physiology of small intestinal epithelial cells in the offspring by shaping their epigenome. Using the transitory colonization system where the animals return to germ-free status ^27^, we asked whether the timing of exposure to microbes or their metabolites would matter for epithelial development and whether there were specific windows pre- or postnatally that induce specific effects on intestinal gene expression and/or chromatin landscapes.

We first carried out RNA sequencing in combination with whole-genome bisulfite sequencing of small intestinal epithelial cells isolated from 21-day-old germ-free C57BL/6 mice. These had been born either to germ-free dams or gestationally colonized dams treated with auxotrophic *E. coli* HA107 ^27^ on intrapartum days 7, 9, 11, and 14. We verified that the transiently colonized dams had returned to germ-free status prior to parturition ^4^, so their offspring remained germ-free throughout the experiment in both experimental groups (Fig. 1A). The protocol was also extended to ask whether there is a window of transient postnatal colonization of the pups themselves that would recapitulate or complement the gestational colonization stimulus, or whether an additional stimulus for active transcription of the preconditioned chromatin landscape is necessary. For this we re-exposed separate groups of germ-free or gestationally primed pups postnatally prior to weaning with the reversible *E. coli* HA107. These experimental groups were termed PnC (postnatal colonization only) and G/PnC (gestational and postnatal colonization) respectively (Fig. 1A).

**Fig. 1.**
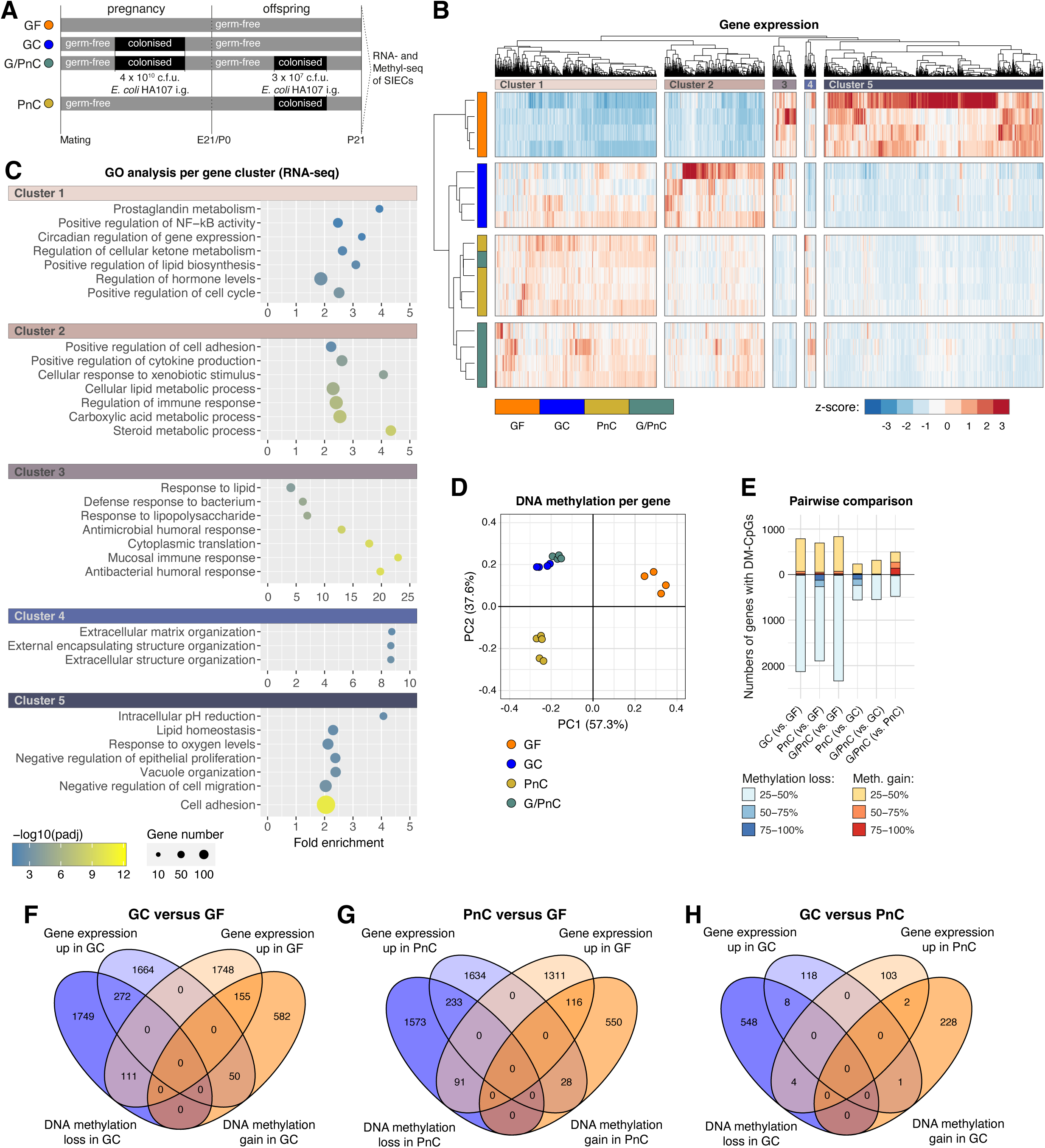
HA107 stimuli during pregnancy and/or postnatally shape gene expression of small intestinal epithelial cell in the offspring partially via DNA methylation. **A.** Germ-free C57BL/6 mice were treated with reversible *E. coli* HA107 during pregnancy (GC - gestationally colonized). Control mice were kept germ-free. Pups born to germ-free or gestationally colonized mice were rechallenged postnatally before weaning (G/PnC - gestational and postnatal colonization or PnC - postnatal colonization). Small intestinal epithelial cells were collected on postnatal day 21 to perform RNA sequencing (B, C) and bisulphite sequencing (D-G) (n = 4 - 5). **B.** Heat map showing differential gene expression between the four groups (fold-change ≥ 2; padj < 0.05). The color scale indicates the relative expression profiles per sample. **C.** GO analysis of the genes in the clusters generated in B. Nodes are colored according to the padj and the size is relative to the number of genes per GO term. **D.** Principal component analysis (PCA) after Pearson correlation based on the differential DNA methylation (≥ 25%; padj < 0.05) landscape restricted to regions ± 20’000 bp to the transcriptional start site (TSS). CpGs located within ±20 kb of annotated transcription start sites were retained, while intergenic CpGs were excluded. For genes represented by multiple CpGs, methylation values were averaged to obtain a gene-level methylation value for each sample. Genes with missing methylation measurements in any of the samples were excluded. In total 416 features were used to generate the PCA plot. **E.** Number of genes with differentially methylated CpGs (DM-CpGs) for each pairwise comparison. Loss and gain are indicative for the first mentioned group relative to the group in brackets. **F-H.** Overlap between significant changes in RNA-seq (fold-change ≥ 2; padj < 0.05) and DNA methylation ((≥ 25%; padj < 0.05) on a gene level between the GC versus GF group (F), the PnC versus GF group (G) and the GC versus PnC group (H) displayed as Venn diagrams.

Transient HA107 exposure strongly affected the gene expression pattern in the offspring regardless of the exposure window, and the four experimental groups formed clusters, all of which were distinct from the transcriptional profile of germ-free mice (Fig. 1B). In some cases, pre- and postnatal colonization (G/PnC) synergized to enhance or abrogate the gene expression pattern observed in germ-free pups. Gene set enrichment analyses showed transcriptional profiles of lipid biosynthesis, response to xenobiotics, and immune mechanisms (clusters 1 and 2) upregulated in pre- or postnatally HA107-exposed compared to the germ-free mice (Fig.1C and fig. S1). In contrast, germ-free mice showed enhanced transcription of genes responsive to lipids, vacuole organization, pH reduction and oxygen levels (clusters 3 and 5), implying delayed epithelial maturation which is likely still adapted for processing maternal milk (Fig. 1C and fig. S2).

To determine whether changes in the abundance of different epithelial subsets could contribute to the observed changes in gene expression we mapped our experimental transcriptomes to single cell database resources that disambiguate different epithelial cell subtypes. Major differences in epithelial cell subtype abundances in germ-free mice compared to the colonized groups were not seen (Fig. S3A-F) indicating that the overall epithelial differentiation program is not disturbed.

In contrast to the presence of transcriptional signals showing normal epithelial subtype development, we did find evidence of altered epithelial proliferation and maturation. Gene set enrichment analysis (GSEA) of differentially expressed genes between GF and GC offspring revealed positive enrichment of the gene sets for ‘G2M checkpoint’ and ‘mitotic spindle’ in the GC groups, emblematic of cell proliferation (Fig. S4A). GSEA also showed positive enrichment in the GF group for ‘myogenesis’, ‘K-Ras signaling down’ and ‘epithelial mesenchymal transition’ (Fig. S4B). Experimental knockdown of *Kras* in the intestinal epithelium triggers a transcriptional portfolio leading to cell-cycle arrest and apoptosis ^28^. ‘Epithelial-to-mesenchymal transition’ in the neonatal intestine reflects activation of developmental programs linked to epithelial plasticity, cytoskeletal remodeling, and growth factor responsiveness ^29,30^. The persistence of these programs in germ-free mice at postnatal day 21 therefore suggests epithelial immaturity in the absence of microbial colonization.

Although the gestationally colonized mothers were germ-free at parturition, the molecular diaspora of their microbial exposure extends through early lactation ^4^. We therefore asked how far these changes are the direct consequence of ongoing microbial exposure compared with potential alterations of the open epigenetic landscape by whole genome bisulfite sequencing. Clustering of the overall DNA methylation landscape per gene locus of the experimental groups revealed that germ-free mice were most distinct from the other groups (Fig. 1D). In addition, in contrast to the transcriptome, postnatal colonization alone in PnC mice did not induce epigenetic changes entirely comparable to those observed in gestationally primed mice (Fig. 1D), indicating that offspring intestinal epithelial cells are strongly shaped by signals originating from the maternal microbiota that cannot be recapitulated through postnatal endogenous microbiota. Pairwise comparisons showed that differences in DNA methylation were the result of a combination of several hundreds of genes with CpGs that had lost or gained methylation and the largest number of CpGs were differentially methylated between the GF and all other groups (Fig. 1E).

A closer look at the GF versus GC group comparison revealed that 2132 genes were affected by methylation loss and 787 genes were affected by methylation gain. Of these, gene ontology (GO) analysis showed methylation gain in genes with metabolic functions, including mitochondrial electron transport, lipoprotein particle clearance, amino acid metabolism and gut morphogenesis (Fig. S5A). Methylation loss was associated with GO terms related to additional metabolic processes and to IL-5 production and actin filament network formation (Fig. S5A), the latter two being indicative of immune activation and intestinal barrier formation, respectively.

To understand to which extent DNA methylation changes induced through the maternal or the early postnatal microbiota would result in altered gene expression at P21, we looked for overlap in significant changes in both read-outs. The Venn diagrams clearly showed that there is substantial overlap between reduced DNA methylation and increased gene expression (272 genes) as well as between increased DNA methylation and decreased gene expression (155 genes) in the GC group compared to the GF group, indicating that the methylation changes induced through maternal microbiota do affect gene expression at the time of weaning (Fig. 1F). The same was true when comparing the PnC group to the GF group (Fig. 1G). Interestingly, the comparison between offspring that was exposed to microbial signals either exclusively during pregnancy (GC) or exclusively post birth (PnC) did show hardly any overlap between DNA methylation changes and an altered transcriptome (Fig. 1H). While this was in line with a low number of differentially expressed genes (126), more than 500 genes still showed methylation changes. Our data thus clearly demonstrate that the epigenome is differentially shaped by gestational versus postnatal colonization, while many of the changes induced in the transcriptome can also be recapitulated by postnatal colonization.

Next, we analyzed the DNA methylation of those genes that were differentially expressed between the different groups as shown in the heatmap in Fig. 1B. We plotted the differential methylation of the genes in an independent clustering per cluster of Fig. 1B and displayed it as five independent heat maps each with additional subclusters labeled “a” to “u” (Fig. S5B). Overall, we observed groups of strongly methylated genes in germ-free mice where the same genes in offspring exposed to HA107 either pre- or postnatally (GC, PnC, and G/PnC) were hypomethylated (Fig. S5B). Previous research indicates that microbiota-dependent effects on DNA methylation precede transcriptomic changes, and the genome gradually loses DNA methylation at several gene loci until weaning ^25^. Our findings show that signals even originating from the maternal microbiota can instruct such a loss of DNA methylation during early life development. In some cases, hypermethylation of 3’ CpG islands was associated with increased gene expression, as demonstrated for genes involved in glycosylation required for intestinal barrier formation ^23^. Overall, DNA methylation and gene expression correlated only partially and only for a subset of genes (subclusters 1c, 1f, 2h, 2m, 5r, 5u in Fig. S5B), implying a complex epigenetic regulation and additional transcriptional factors shaping gene expression. A subset of genes from Clusters 1 and 2 (whose expression was downregulated in GF) represented correlative hypermethylation in germ-free mice. These are involved in saccharide and xenobiotic metabolism, cell cycle control, including transporters and the barrier-forming Claudin 15 (Subcluster 1c, 1f, 2h and 2m in Fig. S5B and Supplementary Data S1). Thus, epithelial cell hypomethylation under colonized conditions can be emblematic of matured epithelial cells, whether these occur through the mother or after birth. The distinction in the DNA methylation pattern in the exclusively postnatally colonized group (PnC) observed in Fig. 1D was mostly absent among the differentially expressed genes of Fig. 1B and S5B. Only a small group of 7 genes within subcluster 1d (Fig. S5B, Supplementary Data S1) showed hypomethylation in the PnC group compared to the GC and G/PnC groups and was thus comparable to germ-free conditions indicating epigenetic integration within the intestinal stem cell compartment in response to microbial stimuli present specifically during gestation. Additionally, the PnC group exhibited a set of genes that appeared rather hypomethylated compared to all other colonized and germ-free controls (subclusters 1a, 2k and 5t; Fig. S5B, and Supplementary Data S1). These genes were associated with stress responses, most probably evoked by the recent postnatal colonization in the absence of preparatory signals delivered by the maternal microbiota, implying that the maternal microbiota during pregnancy confers protection from this stress-induced methylation loss.

Our analysis demonstrates that signals from the maternal microbiota or postnatal colonization induce a large set of non-overlapping methylation changes in small intestinal epithelial cells of the offspring that do not result in significant transcriptomic changes, as well as common hypomethylation in a subset of genes that results in similar gene expression changes in both groups at the time of weaning.

We also verified that microbiota-derived molecules could be transferred in the milk of dams that had been transiently colonized in pregnancy ^4^ through metabolomic analysis of the pups’ stomach contents. While the metabolic content of milk collected form GC dams differed on postnatal day 2, it converges later during development with their germ-free counterparts (Fig. S6). Thus, the effect of the maternal microbial diaspora could also result from the persistence of microbial molecular transfer through lactation. While this was congruent with our previous data ^4^, the current analysis does not allow for tracing of bacterial derived metabolites through isotopic labeling, and thus cannot formally exclude altered host expression in the offspring as causality for the observed metabolic differences in the stomach contents.

### ATAC Seq reveals long-term gene accessibility differences between offspring born to GF or GC dams which partially overlaps with gene expression

Given the limitations of methylation as a marker of current genome accessibility and subsequent gene expression, we prioritized ATAC sequencing of isolated bulk intestinal epithelial cells between the GF and GC groups, since gene expression patterns of GC were generally representative of the PnC and G/PnC groups (see Fig. 1B).

Ordination analysis showed a distinct chromatin landscape between the offspring born to gestationally colonized mothers compared to those born to germ-free control dams (Fig. 2A). A total number of 60,368 consensus peaks (present in at least 3 out of 4 replicates in at least one experimental group) could be called. Of these 23,636 were consensus peaks present exclusively in the GC group, with only 1112 peaks exclusively to the GF group (Fig. 2B). Significant differences in chromatin accessibility between both conditions were assessed performing differential accessibility (DA) analysis on consensus peaks. The genomic distribution of differentially accessible chromatin regions differed strikingly between regions that were differentially more accessible in GC and those that were differential more accessible in GF intestinal epithelial cells. (Fig. 2C). Germ-free IECs exhibited a higher proportion of differentially accessible peaks at transcriptional start sites, consistent with a promoter-centric and developmentally immature chromatin architecture. In contrast, gestationally colonized mice showed an enrichment of differentially accessible peaks within intronic regions, reflecting microbiota-driven activation of distal regulatory elements and maturation of enhancer-based transcriptional programs. This shift from promoter-to enhancer-focused accessibility may highlight the role of early microbial exposure in accelerating postnatal epithelial differentiation and functional specialization.

**Fig. 2.**
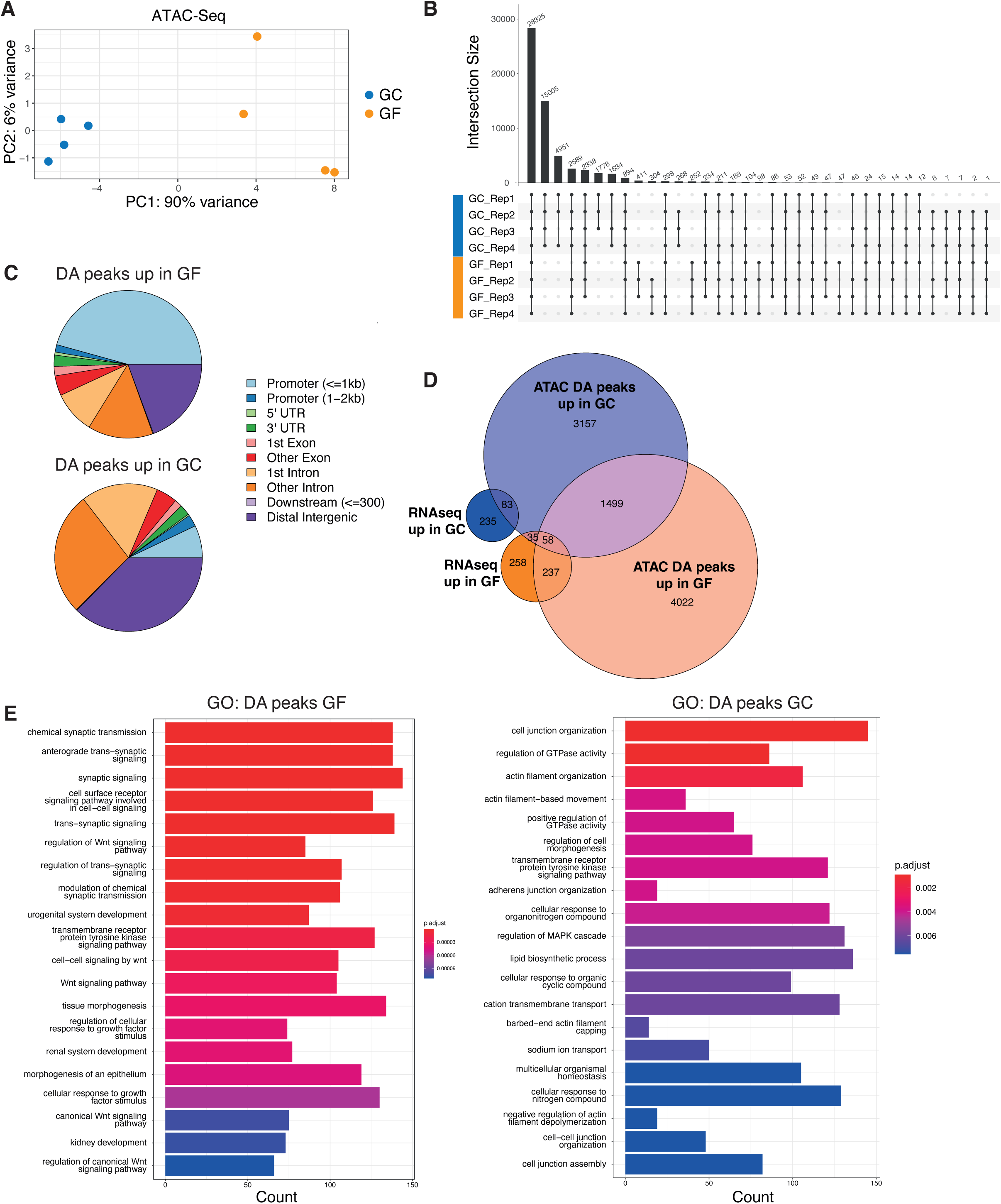
Gestational colonization has a long-term impact on gene accessibility patterns in small intestinal epithelial cells in offspring. Germ-free C57BL/6 mice were treated with reversible *E. coli* HA107 during pregnancy (GC). Control mice were kept germ-free (GF). Small intestinal epithelial cells were collected from offspring on postnatal day 21 and used for Assay for Transposase-Accessible Chromatin (ATAC) sequencing. **A.** Principal component analysis (PCA) of the variance-stabilized count matrix of all filtered consensus peaks (n = 47,049). **B**. Consensus peak distribution of peaks present in at least 3 replicates of either the GF or the GC group. **C.** Genomic distribution of differentially accessible peaks split by peaks significantly more accessible in GF (top chart, n=5281) or in GC (bottom chart, n=7535) offspring. **D**. Venn diagram demonstrating gene overlap between differentially expressed genes in RNA sequencing and genes associated to differentially accessible (DA) chromatin regions in ATAC sequencing results. Up refers to stronger signal in the corresponding experimental condition. Cut-off RNA-Seq: fold-change ≥ 2; padj < 0.05; ATAC-Seq: padj < 0.05. **E**. GO enrichment analysis (Biological Process) of genes associated with DA peaks, splited by left: genes associated with DA peaks more accessible in GF and classified as promoter-peaks; right: genes associated with DA peaks more accessible in GC and classified as intronic-peaks.

We also consolidated the ATAC-Seq data with our RNA Seq data. A proportion of genes significantly increased in expression showed an overlap with differentially accessible chromatin in the same gene region in either of the groups (Fig. 2D). Gene Ontology terms, including organogenesis, neural crest cell migration and differentiation, mesenchymal cell migration, and regulation of growth factor signaling featured in differentially expressed genes with differentially open chromatin in the GF group. Although these terms are not specific to the intestinal epithelium, they reflect conserved embryonic regulatory programs and cellular plasticity pathways that are broadly used during early tissue development ^29^. Their enrichment in germ-free intestinal epithelial cells reflects persistence of an immature, developmentally permissive transcriptional state even at postnatal day 21 (Fig. S4C). GO term biological processes analysis of all gene regions associated with GF-specific peaks at either the promotor or the intron independent of transcriptional changes showed a large proportion of processes being related to Wnt signaling (Fig. 2E) which is also indicative of a less differentiated more stem cell-like nature of the cells ^31^. In contrast, in the GC group-specific peaks, pathways related to the synthesis of junctional complexes and cytoskeletal components were enriched indicative of a tighter, more differentiated mature epithelium, in addition to signals for transport and metabolism of dietary components (Fig. 2E).

We concluded that in contrast to the methylation patterns which are known to have inexact correlation with open chromatin - especially in regard to temporal discordance at enhancers and gene bodies ^32^ - the open chromatin distribution in small intestinal epithelial cells at postnatal day 21 showed a clear maternal microbiota effect that matures the epithelium for functional challenges of postnatal life.

Of these different changes, we next prioritized a detailed examination of epithelial junctional complexes, energy homeostasis, the transcriptional control of maturation and the resultant metabolic phenotype at weaning as functional outcomes of the microbiota-driven epithelial developmental program.

### Maternal exposure during pregnancy or postnatal colonization assists in epithelial maturation via DNA methylation and expression of BLIMP-1 regulated genes

We examined how specific regulators required for replacement of the neonatal by adult epithelial cells were controlled by microbiota-directed epigenetics. BLIMP1, encoded by *Prdm1*, is a master regulator of this differentiation process and acts as a highly-expressed transcriptional repressor in the neonatal intestine ^11,12^. We observed increased *Prdm1* expression in 21-day-old germ-free mice compared to animals from gestationally or postnatally transiently colonized litters (Fig. 3A). *Prdm1* differential expression was matched by distinct transcription patterns of neonatal and adult epithelial signature genes known to respond to the declining levels of BLIMP1 during epithelial transition (Fig. 3B) ^11,12^. Neonatal signature genes tended to be durably expressed in germ-free mice, while adult epithelial signature genes were downregulated compared to HA107-exposed mice (Fig. 3B). Thus, germ-free mice retained an epithelium characteristic of the neonatal period in terms of gene expression, while the DNA methylation indicated a transitional state with intermediate methylation of two neonatal signature genes, *Asl* and *Naga*, whilst adult signature genes remained hypermethylated (Fig. 3C).

**Fig. 3.**
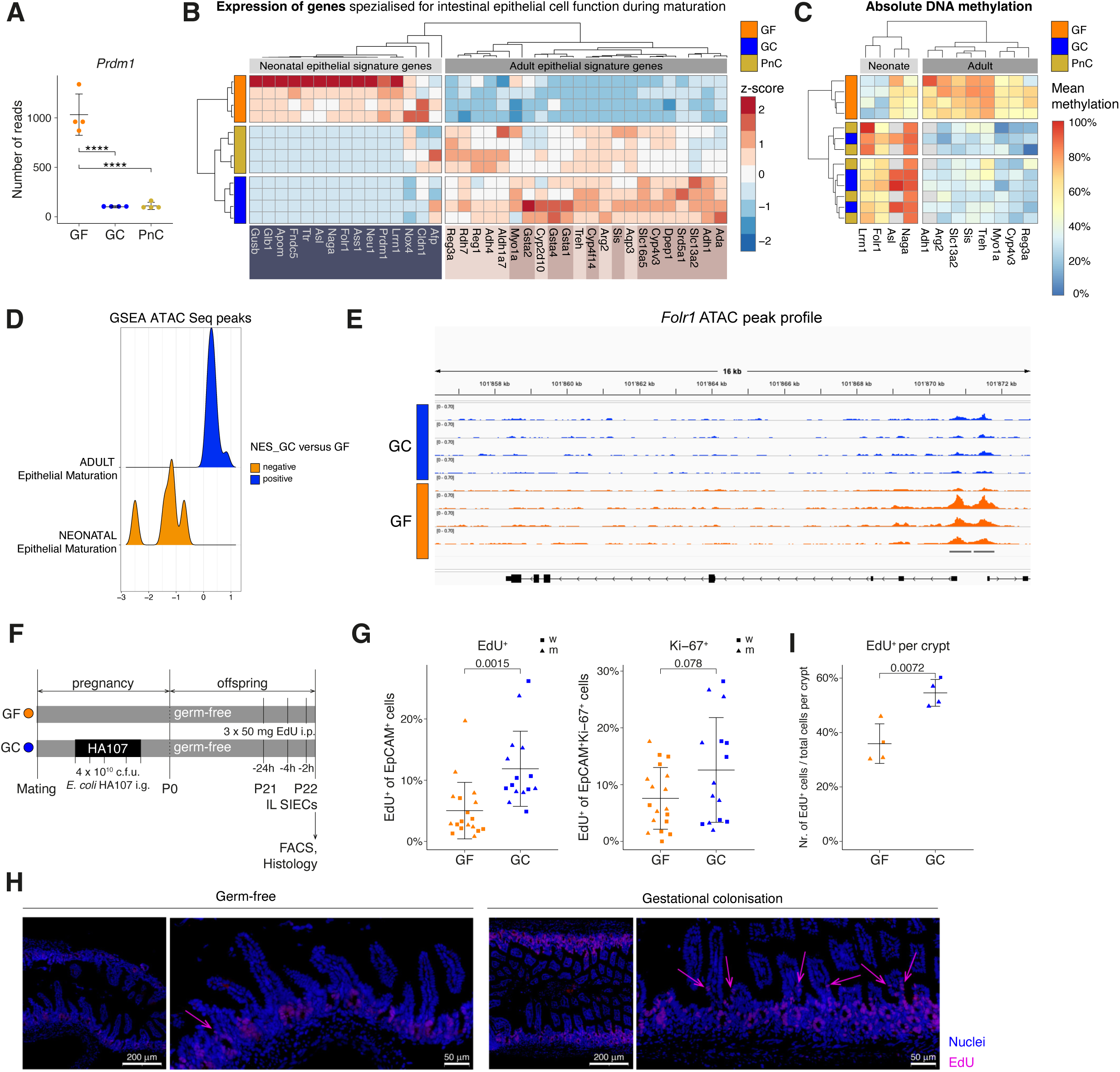
The maternal microbiota contributes to intestinal maturation by suppressing BLIMP-1 thereby mediating epithelial transition. **A.** Normalized reads of *Prdm1* in small intestinal epithelial cells isolated from 21-day-old germ-free (GF) pups, pups born to gestationally colonized dams (GC), and germ-free pups treated with HA107 during weaning (PnC). Asterisks indicate the q-values (**** q ≤ 0.0001), and error bars display the mean ± standard deviation. **B.** Differential gene expression (fold-change ≥ 2; padj < 0.05) of neonatal and adult epithelial signature genes as described by ^11,12^) depicted as heat map. The color scale indicates the relative expression profiles per sample. Genes colors correspond to the clusters in Figure 1. **C.** Heat map showing significant DNA methylation differences (≥ 25%; padj < 0.05). The color scale denotes the mean methylation of all differentially methylated CpGs within ± 20’000 bps relative to the TSS. **D.** Gene Set Enrichment Analysis (GSEA) of differentially accessible chromatin regions at promoters of genes using using the neonatal and adult epithelial signature gene sets. **E.** Genome browser ATAC-Seq peak profiles (CPM normalized) for *Folr1*. **F.** Offspring born to germ-free or gestationally colonized dams were administered three doses of EdU i.p. starting on postnatal day 21. One day later, small intestinal epithelial cells were isolated and EdU incorporation was analyzed with flow cytometry. Cell migration was illustrated in histology of the terminal ileum. Data from two independent experiments were pooled (experiment 1: n GF = 6, n GC = 8; experiment 2: n GF = 12, n GC = 7). **G.** Frequency of EdU^+^ cells in the EpCAM^+^ subset and frequency of EdU^+^ cells among EpCAM^+^Ki-67^+^ cells. The error bars display the mean ± standard deviation. Asterisks indicate p-values with ** p < 0.01. **H.** Representative sections after fluorescent EdU detection. Nuclei were stained in SYBR green (blue) and EdU positive cells were stained in pacific blue (pink). Arrows highlight migrating EdU positive cells. **I.** Quantification of the EdU positive cell number in relation to the total cell number residing in the crypts. The error bars display the mean ± standard deviation. Asterisks indicate p-values with ** p < 0.01.

GSEA of the identified ATAC Seq peaks revealed that the neonatal gene signature showed a positive enrichment score in the germ-free compared to the GC groups (Fig. 3D). This was mainly driven by accessibility changes in *Ass1*, *Folr1*, *Gusb*, *Lrrn1*, *Nox4* as shown representatively for *Folr1* here (Fig. 3E), clearly indicating that silencing of the gene loci of typical neonatal signature genes is accelerated even in mice that had received microbial signals indirectly from the mother.

We next asked when this influence of the microbiota becomes established. In 14-day-old offspring differential BLIMP1-mediated regulation was not found in gestationally colonized pups compared to germ-free mice, as both *Prdm1* expression and its target genes exhibited similar expression levels (Fig. S7A and B), and the DNA methylation was unaffected at this early stage (Fig. S7C). *Prdm1* expression first differed between GF and GC pups at postnatal day 17 (Fig. S7D). Interestingly, by postnatal day 30, expression levels between the two groups had become comparable (Fig. S7D). This suggests that although the program for epithelial transition can be engaged in lowly abundant stem cells by signals originating from the maternal or endogenous microbiota early in life, it may only become relevant for a short duration at the time of weaning. Even without exposure to microbial signals through life, germ-free offspring can eventually transition from the neonatal to the adult epithelium, highlighting an evolutionary hard-wired developmental program for IEC maturation which is fine-tuned by early life microbial signals.

The maturation of the epithelium early in life is accompanied by the acceleration of intestinal epithelial turnover ^11,33^. At weaning, the epithelium undergoes a drastic maturation, including the elongation of the crypt-villus axis and enhanced cell migration ^11,33^. Our transcriptomic analysis has shown that the maternal or postnatal microbiota alters cell cycle regulation without broadly affecting epithelial cell composition (Fig. 1 and Fig. S4) and accelerates BLIMP-1-guided epithelial gene expression changes before weaning (Fig. 3A-C).

To address cell proliferation specifically in the small intestine at weaning, we exposed pups born to germ-free and gestationally colonized dams to three doses of 5-ethynyl-2’-deoxyuridine (EdU) (Fig. 3F). One day after EdU injection, small intestinal epithelial cells of the ileum were isolated and the frequency of EdU^+^ cells among EpCAM^+^ epithelial cells was assessed by flow cytometry (Fig. 3G and fig. S7E). The gestationally colonized group displayed a higher frequency of EdU^+^ cells among the EpCAM^+^ subset (Fig. 3G), which was confirmed in histology and demonstrated enhanced cellular trafficking towards the villus tip (Fig. 3H and I). However, the frequency of cycling EpCAM^+^Ki-67^+^ cells remained stable (Fig. 3G). Limited expansion in the proliferative zone, as determined by the unaffected EpCAM^+^Ki-67^+^ cell pool along with enhanced migration of EdU^+^ cells relative to germ-free offspring, indicates less sustained cellular cycling with accelerated turnover upon gestational colonization. Given the observed alterations in the epigenetic landscape, gene transcription and EdU incorporation, we propose that the maternal microbiota primes intestinal epithelial cells to regulate epithelial turnover and to accelerate epithelial maturation along with increased cell migration at weaning. Although germ-free mice catch up and eventually transition from the neonatal to the adult epithelium, early life microbial signals fine-tune the timing of this transition to occur at the time of weaning, a period characterized by major environmental pressures.

### The maternal or postnatal microbiota drives maturation of junctional complexes in the offspring via transcriptional regulation

An important feature of postnatal epithelial differentiation is the development of intercellular junctional complexes. Mouse tight junction proteins follow specific postnatal developmental waves with pore-forming claudins being highly expressed early in life, thereby increasing intestinal permeability, while barrier-forming molecules are more abundant during adulthood ^14–16^. In line with ATAC results for open chromatin for adhesion GO-terms, we confirmed that genes for cell adhesion molecules were generally upregulated and accessible in the GC group, whereas claudin genes were repressed upon HA107 stimuli (Fig. 1B, fig. S1, S2). Mapping these developmental and functional characteristics of claudins ^14–16^ and their regulatory circuits ^34–38^ with our dataset, claudin genes selectively expressed during the neonatal period - including *Cldn2* and *Cldn8* - were downregulated by gestational or postnatal HA107 treatment, whereas transcription of the adult claudins *Cldn3* and *Cldn15*, and the barrier-forming junctional complex Zonulin-1 (*Tjp1*), were enhanced (Fig. 4A, B and fig. S8A and B).

**Fig. 4.**
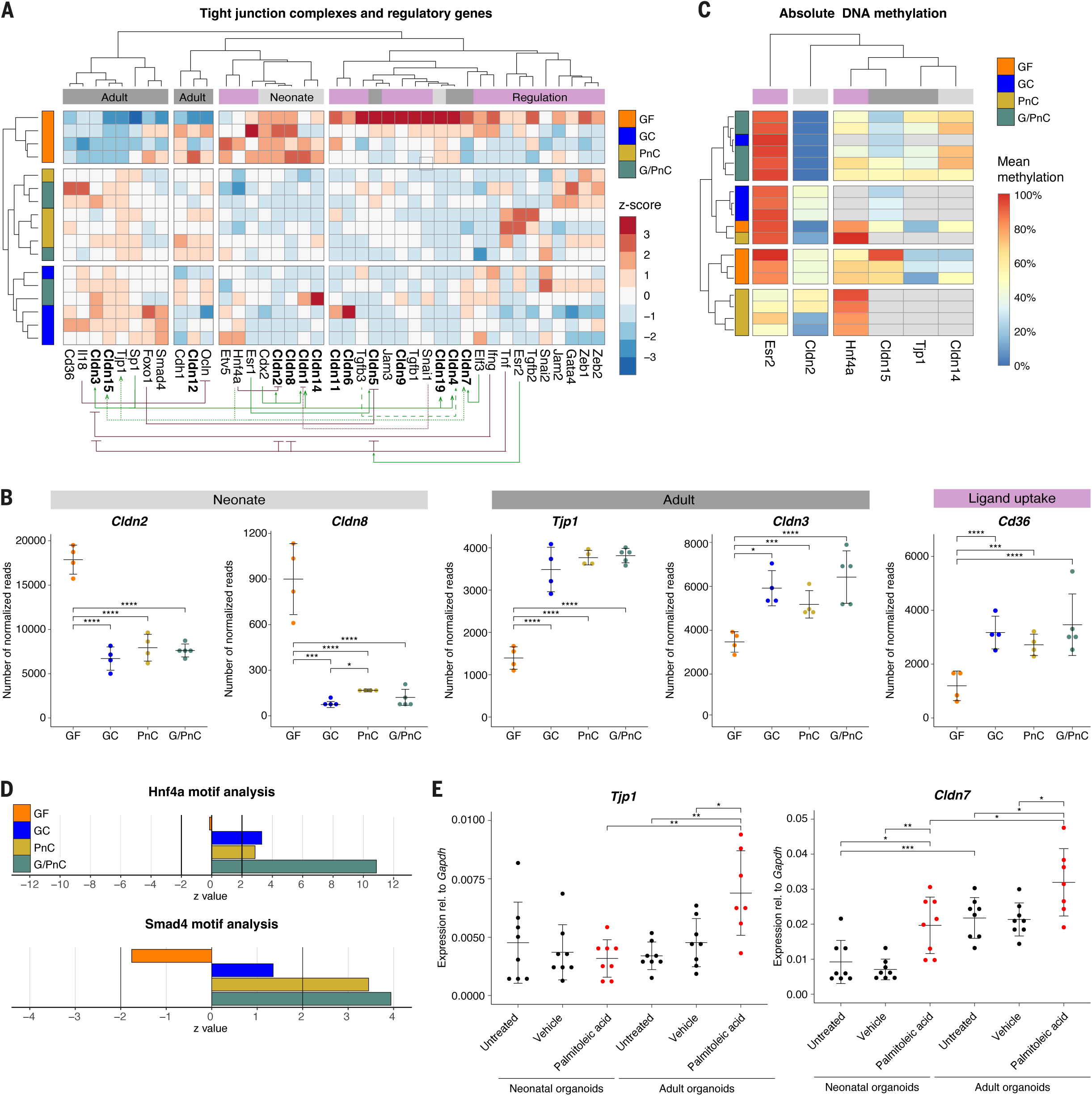
Gestational colonization facilitates the development of barrier-forming tight junctions by regulating gene expression. **A.** Heat map illustrating differential gene expression (fold-change ≥ 2; padj < 0.05) of tight junction complexes and associated transcription factors with unsupervised clustering categorized based on neonatal or adult gene signatures ^14–16^. Green arrows signify transcription factors that promote expression, while red arrows denote those involved in negative regulatory interactions ^34–38^. **B.** Number of normalized reads in the RNA sequencing analysis of genes encoding various tight junction molecules and the fatty acid transporter *Cd36*. Unpaired parametric student’s t-test was performed. The error bars display the mean ± standard deviation. Asterisks indicate q-values with * q < 0.05, *** q ≤ 0.001 and **** q ≤ 0.0001. **C.** Heat map with the average DNA methylation levels of genes showing significant methylation differences (≥ 25%; padj < 0.05) linked to tight junctions. Grey defines no significant methylation difference in any CpG. **D.** Motif analysis assessed with the RNA sequencing data quantified as z-value, which corresponds to the number of standard deviations that the computational motif’s activity is away from zero. **E.** Expression of *Tjp1* and *Cldn7* relative to *Gapdh* in neonatal or adult small intestinal organoids treated with the HNF4a ligand palmitoleic acid. The error bars display the mean ± standard deviation. Asterisks indicate p-values with * p < 0.05, ** p ≤ 0.01, *** p ≤ 0.001 and **** p ≤ 0.0001.

Since the DNA methylation and ATAC accessibility pattern of tight junctional genes did not show conclusive adaptations by gestational or postnatal exposure (Fig. 4C), we hypothesized an overlay of direct metabolite dependent regulatory mechanisms. Gene expression of claudins including *Cldn2*, *Cldn7* and *Tjp1* is tightly regulated by the ligand-activated hepatocyte nuclear factor 4 alpha (HNF4a) ^38^. Although *Hnf4a* gene expression remained unchanged in our dataset (Fig. S8C), the genes containing the binding motif of HNF4a or its interaction partner SMAD4, in addition to the CD36 gene expressing the scavenger receptor for import of HNF4a ligands, were significantly more expressed in mice after direct or indirect HA107 exposure (Fig. 4B and D). To test whether expression of these junctional genes could be phenocopied by authentic HNF4a ligands, we supplemented the media of neonatal and adult murine small intestinal organoids with the HNF4a ligand palmitoleic acid. Adult-specific junctional transcripts known targets of HNF4a, such as *Cldn7* and *Tjp1*, were significantly increased in adult organoids receiving palmitoleic acid. In organoids derived from 14-day-old pups, palmitoleic acid supplementation was sufficient to increase the expression of *Cldn7* (Fig. 4E).

We concluded that the postnatal developmental dynamics of epithelial junctional development is shaped both by the maternal and postnatal drivers of the epigenetic landscape and by the broadening molecular exposure as the young animals become weaned from lactation.

### Gestational colonization improves energy homeostasis in the neonatal intestine by modulating aerobic respiration

We have shown that bacterial stimuli during early life can stimulate cell cycle processes, tight junctional remodelling and epithelial restructuring during the suckling to weaning period. There was upregulation of genes associated with glycolysis and mitochondrial biogenesis in the offspring born to gestationally colonized dams (Fig. S9). Nuclear genes driving mitochondrial biogenesis curated according to Popov et al. ^39^ and Villena et al. ^40^ also exhibited significantly increased expression following gestational colonization or postnatal colonization (Fig. 5A). The key regulator of mitochondrial biogenesis, peroxisome proliferator-activated receptor-g coactivator (PGC-1a, encoded by *Ppargc1a* and *Ppargc1b*) and co-activators involved in the post-transcriptional activation of PGC-1a, such as as *Sirt1* were significantly upregulated in GC and PnC pups (Fig. 5A and B). The interpretation that early life signals from the microbiota drive developmental processes involving increased mitochondrial biogenesis was supported by poor induction of respiratory chain and ATPase genes in the germ-free group (Fig. S8D). Interestingly, structural genes of the mitochondria were mostly enhanced in the GC group (mttRNA and mtrRNA) whereas mt mRNAs were more abundant in the PnC group (Fig. S8D) suggesting a nuanced or time-dependent regulation of mitochondrial transcription depending on direct exposure to microbes, or indirect exposure to microbial metabolites from the maternal microbiota.

**Fig. 5.**
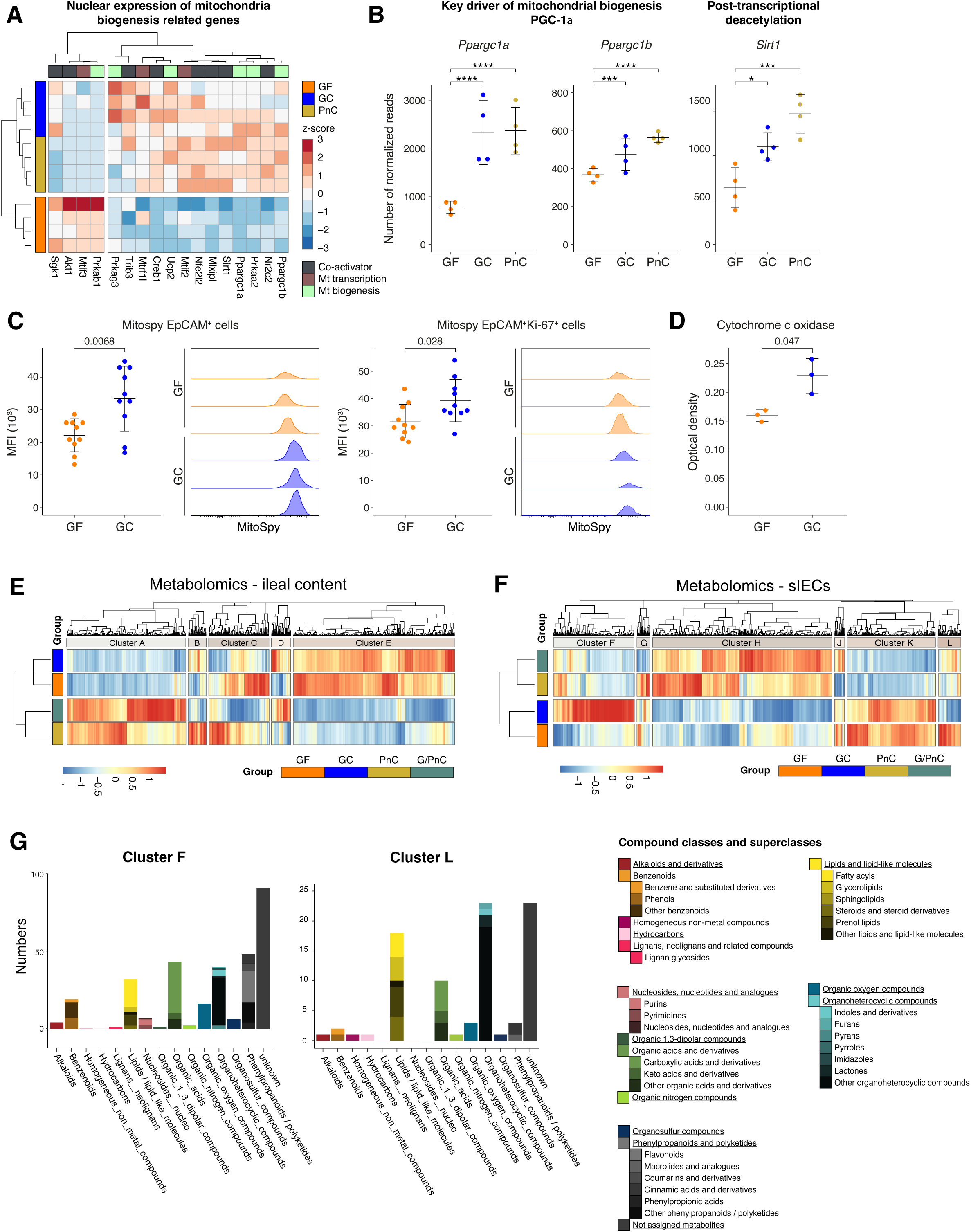
Maternal colonization changes metabolic programs in offspring sIECs at weaning. **A.** Heat map including nuclear genes with significant differential expression between at least one pairwise comparison (fold-change ≥ 0.5; padj < 0.05) involved in mitochondria biogenesis ^39,40^. The color scale illustrating the relative expression profile per sample. **B.** Number of normalized reads of *Ppargc1a*, *Ppargc1b* and *Sirt1*. The error bars display the mean ± standard deviation. Asterisks indicate q-values wit * q < 0.05, *** q ≤ 0.001 and **** q ≤ 0.0001. **C.** Median fluorescence intensity (MFI) and representative histograms of MitoSpy orange CMTMRos in EpCAM^+^ and EpCAM^+^Ki-67^+^ cells isolated from the small intestine of germ-free pups (GF) or pups born to gestationally colonized dams (GC). Data were reproduced in two additional experiments. The error bars display the mean ± standard deviation. Asterisks indicate p-values with * p < 0.05 and ** p < 0.01. **D.** Quantification of the optical density in the cytochrome c oxidase staining calculated from the grey value of 10 villi per section. The error bars display the mean ± standard deviation. Asterisks indicate the p-value with * p < 0.05. **E-G**. Untargeted metabolome analysis of ileal content and small intestinal epithelial cells collected from 21 day-old offspring in the GF, GC, PnC or G/PnC groups. **E**. Differentially abundant metabolites in ileal content are plotted in a heat map. The color scale indicated the relative abundance of metabolites per sample. **F.** Differentially abundant metabolites in small intestinal epithelial cells are plotted in a heat map. The color scale indicated the relative abundance of metabolites per sample. **G.** Distribution of compound classes and superclasses visualized as stacked bar plots for the clusters F and L. See also data in Supplementary Data S2.

We next questioned whether this effect represents crypt-villus partitioning, since mitochondrial function has a crucial role in maintaining epithelial stemness and regulating proliferation in the transit amplifying zone ^41^. We assessed mitochondrial membrane potentials using MitoSpy™ orange chloromethyl-tetramethylrosamine (CMTMRos), which accumulates in the negatively charged space of the mitochondria. Epithelial cells marked with EpCAM and also its cycling subset (EpCAM^+^Ki-67^+^) in pups born to dams exposed to HA107 during pregnancy significantly incorporated more dye into the inner membrane, indicating a negative membrane potential and thus polarized mitochondria (Fig. 5C). In contrast, germ-free mice displayed less polarized mitochondria, likely due to less respiratory chain activity (Fig. 5C). Histochemical cytochrome c oxidase staining in intestinal sections showed activity largely in the epithelial cells, which was quantitatively greater in pups born to dams treated with HA107 compared to germ-free mice (Fig. 5D and Fig. S8E).

These data imply that microbial signals during the prenatal window, via colonization of the mother during pregnancy, facilitate postnatal epithelial energetic development.

### The maternal microbiota programs the metabolic phenotype of the small intestinal epithelium at weaning

Having shown that signals originating from the maternal microbiota affect the molecular regulation to adjust epithelial homeostasis and energy metabolism during early life development, we asked whether early life exposure to microbiota resulted in altered fucntional xenobiotic metabolism after weaning. We therefore analyzed the intestinal contents and tissue metabolome after germ-free or GC pups had been weaned onto chow at postnatal day 21.

The most obvious difference was observed between mice with and without postnatal colonization which may be indicative of remaining bacterial metabolites in the intestine (Fig. 6A). Whereas stomach and ileal content metabolomes of GF and GC offspring were comparable, as metabolites from the microbial exposure of the dams four weeks previously had been cleared (Fig. 5E and fig. S6), small intestinal epithelial cells of mice that were exposed to microbiota in utero and weaned on chow diet exhibited an altered metabolite composition compared to their GF control group (Fig. 5F).

Specifically, maternal exposure to HA107 during the prenatal window increased the levels of carboxylic acids, benzenoids and phenylpropanoid xenobiotic metabolites in intestinal epithelial cells (Clusters F, Fig. 5F and G). In contrast, germ-free mice were enriched in lipids (Cluster l, Fig. 5F and G) but displayed fewer organic acids and derivatives and higher levels of arginine (Supplementary Data S2), an amino acid essential early in life, the uptake of which is under the regulation of BLIMP-1. This alines with our sequencing analysis, showing that germ-free mice exhibited higher expression of genes associated with lipid homeostasis and vacuole organization important for lipid uptake before weaning (Fig. 1C and fig. S10). We concluded that the maternal microbiota guides functional epithelial maturation during weaning to adapt to the changing diet from milk to solid food, the latter being contaminated with xenobiotics.

## DISCUSSION

Our model of complete reversibility of colonization during defined early-life time windows uniquely allows the effects of maternal microbiota during pregnancy from later endogenous colonization to be distinguished ^4^. We have shown that commensal stimuli present during early life windows drive gene expression changes at the time of weaning emblematic of intestinal development, including the BLIMP-1 guided neonatal-to-adult epithelial transition, “gut closure”, mitochondria biogenesis and metabolic adaptations at weaning. These effects can even be driven solely by transient maternal colonisation in utero, when the offspring are born and remain germ-free after birth, so they have only ever been indirectly exposed to the mother’s microbial metabolites. The endogenous microbiota of the neonate postnatally can phenocopy many but not all transcriptomic changes of transitory maternal colonisation, highlighting that different early life windows likely allow for precise and time-dependent modulation by environmental factors, including microbial signals.

Some of the observed changes in gene expression could be correlated to an altered chromatin landscape, suggesting epigenetic regulation as one possible mechanism. Interestingly, only a subset of the changes observed in gene expression overlapped with an altered genome accessibilty and vice versa. While maternal microbiota-induced transcriptomic changes could to a large extend be recapitulated by postnatal colonization, the methylation landscape differed tremendously between mice that had been either exclusively exposed to maternal micorbiota or to postantal endogenous microbiota. Postnatal colonization alone resulted in hypomethylation of stress-related genes compared to the influence of gestational colonization or combined gestation and postnatal colonization.

The observation that transcriptomic differences were minor between gestationally exposed and postnatally exposed groups despite the large differences in the epigenetic landscape may be explained by the nature of our experimental model in which the offspring artificially stays germ-free until weaning. These differences in the epigenome may result in more prominent gene expression changes in mice that are exposed to a maturing intestinal microbiota or pathogens during early or later life.

Epigenetic modification via DNA methylation and offering better gene loci accessibility can contribute to these changes. The maternal microbiota specifically targeted and fine-tuned the timing of BLIMP-1-guided epithelial transformation. This key regulator for replacing the immature neonatal epithelium, encoded by *Prdm1,* is highly expressed in the neonatal intestine and gradually decreases until postnatal day 21 ^11,12^. The DNA methylome and ATAC peak profile of *Prdm1* target genes reflected gene transcription patterns, with neonatal signature genes being methylated (inaccesible) and adult signature genes being hypomethylated in the gestational colonization group compared to the germ-free group.

Previous studies have shown that microbial colonization can elicit substantial transcriptional responses in the intestinal epithelium with comparatively limited changes in chromatin accessibility, while identifying histone modifications as an important microbiota-responsive regulatory layer ^9,10,42^. Our findings are not incompatible with these observations but extend them to a distinct developmental context: microbial signals encountered during gestation and early postnatal life coincide with establishment of the epithelial regulatory landscape and the neonatal-to-adult transition. The persistence of distinct chromatin and transcriptional states according to the timing of microbial exposure suggests that developmental history contributes to epithelial regulatory state at weaning. The presence of a continuous endogenous microbiota may add an additional regulatory layer altering gene expression which may be less dependent on modifications on the chromatin level.

Although *E. coli* is an early-life colonizer, it does not recapitulate the full functional breadth of a complex maternal or neonatal microbiota ^43^. Microbial metabolites, including short-chain fatty acids, folate and polyphenol derivatives, are key modulators of the host chromatin landscape, and their production varies substantially across taxa ^44^. *E. coli* does contribute to this pool through *de novo* folate synthesis, polyphenol biotransformation, and acetate production ^45,46^. However, the mechanistic effects of microbial metabolites on the chromatin landscape varies according to the microbial and dietary environment as exemplified in the context of obesity where the epigenetic methylation and histone modification landscape is also responsive to taxon compositional changes ^10,42^. Our objective here was to use a single organism in which we could control the colonisation timing and duration, although this still allowed a heterogenous molecular drive of a large series of different biochemical stimuli for epigenetic alterations, including DNMTs and histone modifying enzymes, whose individual disambiguation is beyond our present resolution.

We previously demonstrated that microbial ligands originating from the maternal microbiota are transferred to the fetus, or the neonate via breast milk, driving maturation and maintenance of innate leukocytes in the intestinal lamina propria ^4^. Our new findings expand on this by showing that gestational colonization further alters the epigenome of intestinal epithelial cells supporting their maturation, making the offspring more adapted and resilient to external clues, such as exposure to commensals, infections, and the dietary switch at weaning.The portfolio of maternal and postnatal microbiota-driven effects extends across epithelial junctional complexes, energy homeostasis, the transcriptional control of maturation and the resultant metabolic phenotype at weaning. Not all of these are driven by chromatin remodelling and our findings propose additional microbiota-dependent effects on direct transcriptional regulation of gene expression and intestinal function, potentially through microbial products or secondary pathways involving immune regulation ^5,16,38,47–49^.

Our study shows the pervasive complementary effects of both the maternal and postnatal microbiota and their metabolite diaspora as an important component of the maternal-fetal-neonatal interface that drives postnatal intestinal development and matures the postnatal intestinal function.

### Limitations of the study

Our study has several limitations. First, reversible gestational colonization with a single *E. coli* strain provides precise temporal control over maternal microbial exposure but cannot recapitulate the taxonomic and metabolic complexity of a diverse maternal microbiota, despite the broad repertoire of metabolites produced by *E. coli*. Second, although our findings demonstrate sustained effects of maternal microbial exposure on offspring epithelial development, we did not resolve the route by which these signals are transmitted. Our current and previous observations are consistent with both transplacental exposure and postnatal transfer of microbiota-derived metabolites through breast milk; experimentally disentangling these routes, for example by cross-fostering, would require substantially expanded animal cohorts and introduce additional stress-associated variables, conflicting with efforts to minimize animal use under the 3R principles. Moreover, while we comprehensively characterized the functional consequences of early microbial exposure for epithelial maturation, barrier development, and metabolic capacity, the specific microbial metabolites responsible and the persistence of these effects beyond weaning remain to be determined. Defining the molecular mediators, routes of mother-to-offspring transmission, and long-term consequences of this developmental programming will therefore be important directions for future studies. Finally, our chromatin analyses focused on accessibility and therefore capture only one layer of epigenetic regulation; microbiota-dependent histone modifications and other chromatin-associated mechanisms may additionally contribute to the transcriptional phenotypes observed.

## MATERIAL AND METHODS

### Mice

Germ-free, gestationally treated dams and gnotobiotic C57BL/6(J) mice were bred and maintained in flexible-film isolators at the Clean Mouse Facility (CMF) of the University of Bern and the Genaxen Foundation in Switzerland. Axenic status was routinely monitored in bedding, food and animal tissues by aerobic, anaerobic cultures and gram stainings. All experiments were performed in accordance with Swiss Federal and Cantonal regulations under license agreements: BE104/20 and BE132/2023.

### Reversible colonization

Pregnant dams were administered 10^10^ c.f.u. *E. coli* HA107 by intragastric gavage on gestational days 7, 9, 11 and 14 determined by vaginal plug observation after mating. Feces were collected three days after the last treatment and tested with liquid amplification and solid media culture. For HA107 rechallenge experiments, age-matched pups born to gestationally treated dams and germ-free mice were administered 10^7^ c.f.u. *E. coli* HA107 on postnatal days 14, 16 and 19 by intragastric gavage. Germ-free status of rechallenged P21 pups was confirmed with SYTOX green staining of colon content.

### Bacterial culture

*E. coli* HA107 ^27^ was cultured at 37°C, 200 rpm overnight in LB medium supplemented with 100 µg/ml m-DAP and 400 µg/ml D-Ala. Bacteria were centrifuged for 10 min at 4000 rcf, washed with sterile PBS and reconstituted in 500 µl sterile PBS per adult mouse or 200 µl per neonate. Auxotrophic status was confirmed by plate culture on LB without supplements and bacterial load per gavage was assessed on LB agar containing 50 µg/ml m-DAP and 200 µg/ml D-Ala.

### Small intestinal epithelial cell isolation

The ileum was removed from the euthanized mouse, opened and cleaned to remove intestinal contents and mesenteric fat. After washing in DPBS, epithelial cells were separated by incubating the tissue in dissociation buffer (DPBS, 5 mM EDTA, 10 mM HEPES) at 37°C on a magnetic stirrer for two periods of 10 min. Epithelial cells from both fractions were passed through a 100 µm cell strainer and collected in IMDM with 10% FCS. The cells were pelleted (2000 rcf, 10 min, 4°C) and resuspended in IMDM/FCS. Intraepithelial lymphocytes were depleted by Percoll separation. For this, cells were overlayed onto 40% Percoll (GE Healthcare Bio-Sciences) in pure IMDM, centrifuged at 670 rcf for 30 min (RT), acceleration at 3 and break 1 (Eppendorf Centrifuge 5810). The intermediate phase containing the epithelial cells was collected in IMDM/FCS. Percoll separation was omitted for flow cytometry. After washing in DPBS, pelleted epithelial cells were resuspended in the respective buffer or snap-frozen in liquid nitrogen for subsequent experiments.

### RNA extraction

Small intestinal epithelial cells were collected in Trizol. Molecular separation was achieved with chloroform separation upon centrifugation (15 min, 12’000 rcf, 4°C). The aqueous phase was washed with ice-cold isopropanol (12’000 rcf, 10 min, 4°C) followed by precipitation in ice-cold 75% ethanol (7500 rcf, 5 min, 4°C). The RNA was air-dried, resuspended in DNase/RNase-free distilled water and purified with the RNAse-free DNAse kit (Qiagen) and the RNAeasy MinElute kit (Qiagen) according to the manufacturer’s instructions. Concentration and quality were determined with the Bioanalyzer 2100 (Agilent).

### cDNA synthesis and real-time PCR

RNA (1-5 µg) was reverse transcribed using Superscript III reverse transcriptase (Invitrogen) and random hexamers (Promega) according to the manufacturer’s protocol on a C1000 touch thermal cycler (Biorad). The relative gene expression was measured by real-time PCR with 8 - 25 ng of cDNA per reaction, gene-specific primers (Supplementary Material) and the QuantiNova SYBR Green PCR kit (Qiagen). All reactions were run in triplicate for 40 cycles on a CFX opus real-time PCR system (Biorad). Expression of the gene of interest (GOI) was normalized according to the cycle threshold (Ct) of the housekeeping gene *Gapdh* using the formula 2^-ΔCt^ (ΔCt = CtGOI-CtGapdh).

Primer sequences:

*Gapdh*_for: CATCAAGAAGGTGGTGAAGC,

*Gapdh*_rev: CCTGTTGCTGTAGCCGTATT,

*Tjp1*_for: GAGCGGGCTACCTTACTGAAC,

*Tjp1*_rev: GTCATCTCTTTCCGAGGCATTAG,

*Cldn7*_for: ACCAACTGCGTACAAGACGAG,

*Cldn7*_rev: CAGAGCCGCCAACAGGAAA

### RNA sequencing and analysis

Libraries were prepared with the TruSeq RNA sample preparation v2 kit (Illumina) and sequenced on an Illumina HiSeq 3000 on 150 bp paired-end mode. The quality of the reads was assessed using fastqc v.0.11.5. The reads were mapped to the GRCm38 reference genome with Hisat2 v.2.1.0 ^50^. FeatureCounts from Subread v.1.6.0 was used to count the number of reads overlapping with each gene, as specified in the Ensembl annotation (release 101) ^51^. The Bioconductor package DESeq2 v.1.24.0 in R v.3.6.1 was used to test for differential gene expression between experimental groups with read counts normalized over all samples ^52^. GO terms significantly enriched for differentially expressed genes were identified with topGO v.2.38.1 in R v.3.6.1. All tests were repeated using different combinations of algorithms (weight01 or classic) and test statistic (Fisher or Kolmogorov-Smirnov) to assess the robustness of the results. To generate heatmaps, all genes differentially expressed in any pairwise comparison between groups (log2-fold-change >1; adjusted p-value <0.05) were plotted. Z-scoring was performed gene wise across samples and the heatmap was generated using the pheatmap function in R. The Cytoscape software v.3.9.0 was used to visualize genome networks ^53,54^. The functional relationship in the complete network context was predicted by performing a STRING enrichment analysis using the integrated stringApp v.1.6.0 with an enrichment significance threshold set to the default value 0.05 ^55^. Most significant and relevant GO terms (FDR ≤ 0.05) were selected according to Markov clustering performed by clusterMaker2 v.1.3.1 implemented in the Cytoscape software to cluster according to STRING networks ^56^. Motif enrichment analysis was performed with the Integrated System for Motif Activity Response Analysis (ISMARA) ^57^. *In silico* analysis of cellular proportions was performed with the online analytical tool CIBERSORTx ^58^ based on published single-cell data of murine small intestinal epithelial cells ^59^.

### Whole genome bisulphite sequencing

For bisulphite sequencing, small intestinal epithelial cells were snap-frozen in liquid nitrogen and resuspended in DPBS. The DNA was purified with the Epicentre Masterpure DNA Purification Kit (Lucigen) according to the manufacturer’s instructions. DNA (50 ng) was sheared to a length of 350 bps on a E220evolution focused-ultrasonicator (Covaris). Bisulphite conversion was performed using the EZ DNA Methylation-Lightning Kit (Zymo Research), following the manufacturer’s protocol. Libraries were prepared with the TruSeq DNA Methylation kit (Illumina) and sequenced on an S4 flow cell with an Illumina HiSeq 3000 on 150 bp paired-end mode. Initial processing and quality control was performed using the nf-core methylseq pipeline v.1.4. To map the reads to a modified GRCm38 reference genome and obtain methylation calls Bismark v.0.22.2 was used ^60,61^. This modified reference was produced by changing selected positions in the GRCm38 sequence to match the sequence observed in whole genome resequencing data from this mouse strain. Downstream analyses were run using our own pipeline implemented in Snakemake. Specifically, methylKit v.1.14.2 in R v.4.0.2 was used to identify differentially methylated CpGs ^62^. Metilene v.0.2-8 ^63^ and dmrseq v.1.8.0 ^64^ in R v.4.0.2 were used to identify differentially methylated regions (DMRs) and created the union of DMRs from the two tools. Individual CpGs and DMRs were annotated using the annotatePeaks.pl script from Homer v.4.11 ^65^.

### ATAC sequencing

To profile open chromatin regions in cells using ATAC-Seq, 60’000-90’000 cells of high quality and viability were prepared. ATAC-Seq libraries were made using an Active Motif ATAC-Seq Kit (Active Motif, 53150) following their protocol (revision version B11) for cell sample preparation, tagmentation, purification and PCR Amplification of tagmented DNA. An ATAC-Seq Spike-In Control (Active Motif, 53154) was used wherein 10’000 spike-in nuclei were added to the sample cells after sample cell counting but before centrifugation to pellet the sample cells. The quantity and length of the generated DNA libraries was evaluated using a Thermo Fisher Scientific Qubit 4.0 fluorometer with the Qubit dsDNA HS Assay Kit (Thermo Fisher Scientific, Q32854) and an Agilent Fragment Analyzer (Agilent) with a HS NGS Fragment Kit (Agilent, DNF-474), respectively. The libraries were pooled equimolarly and sequenced in paired-end mode using NextSeq 1000/2000 P2 Reagents v3 (200 cycles; Illumina, 20046812) on an Illumina NextSeq 1000 instrument. The quality of the sequencing runs was assessed using the Illumina Sequencing Analysis Viewer (Illumina version 2.5.12) and all base call files were demultiplexed and converted into FASTQ files using Illumina bcl2fastq conversion software v2.20. All steps post cell counting were performed at the Next Generation Sequencing Platform, University of Bern.

Raw ATAC-seq data were processed using the nf-core/atacseq pipeline (v2.1.2) ^66^ using the following specified settings, and default otherwise: reads were aligned to *Mus musculus* reference genome (GRCm38), read length specified for MACS2 was 75, biological replicates were treated as independent samples, peak calling was performed in “narrowPeak” mode, and peaks were retained if they were detected in at least three biological replicates (≥3). We refer to these peaks as consensus peaks.

Downstream analysis and visualization of consensus peaks (60368) were performed in R (v4.2.1). For differential accessibility analysis, low-signal peaks were filtered out by retaining only those with ≥50 reads in at least two samples (leading to a total of 47049 filtered consensus peaks). DA analysis was performed on the filtered count matrix using DEseq2 R package (v1.38.3) ^67^. Peaks with adjusted *p* < 0.01 and absolute log2 fold change > 0 were considered significantly differentially accessible. Peak annotation based on genomic feature overlap was performed using the ChIPseeker R package (v1.34.1) ^68^. Gene annotations were derived from the TxDb.Mmusculus.UCSC.mm10.knownGene (v3.10.0). Promoters were defined as −2000 bp to +500 bp from the transcription start site.

Peaks annotated as overlapping gene promoters (promoter-peaks, n=14846) were selected to perform gene ontology (GO) enrichment analysis and gene set enrichment analysis (GSEA) based on the overlapping genes. GO enrichment analysis was then performed independently for the genes associated with DA promoter-peaks upregulated in each group using the GOenrich() function from the clusterProfiler R package (v4.6.2) ^69^, with adjusted p-value cutoff of 0.01. The background universe consisted of all promoter-peaks identified in the initial peak annotation. The same logic was applied to perform GO enrichment analysis on peaks overlapping introns (intronic-peaks, n=31768). Gene set enrichment analysis (GSEA) was performed using clusterProfiler R package on custom lists of features. If more than one peak overlapped a gene promoter region, the mean fold change of the peaks was taken for GSEA analysis.

RNAseq and ATACseq data were integrated at gene level, by considering DE genes from RNAseq and associated genes to DA peaks from ATACseq. To further perform GO enrichment analysis on the overlap of both technologies, genes exclusively overlapping RNAseq and ATACseq in a up regulation fashion were considered for the analysis.

### EdU cell proliferation assay

EdU incorporation assays were performed using the Click-iT™ Plus EdU Pacific Blue™ Flow Cytometry Kit according to the manufacturer’s guidelines (ThermoFisher). Briefly, mice were administered 50 µl 1 mg/ml EdU solution i.p. 24 h, 4 h and 2 h before small intestinal epithelial cell isolation. Dead cells were stained with fixable viability dye (eBioscience) for 20 min on ice. After washing in PBS, extracellular staining of Epcam (Biolegend) was performed for 15 min on ice and subsequently washed twice with PBS. Cells were fixed for 15 min with 100 µl Click-iT™ fixative. After washing twice with PBS, cells were resuspended in 50 µl 1X Click-iT™ fixative saponin-based permeabilization and wash reagent and incubated for 15 min. Click-iT™ Plus reaction cocktail was mixed according to the manufacturer’s instructions, with an antibody concentration of 1:200, added to the cells, and incubated twice for 30 min at RT. Intracellular staining of Ki-67 (eBioscience) was performed overnight. Samples were acquired using an LSRFortessa (BD Biosciences) and data were analysed using FlowJo™ v.10.8.1 (BD Biosciences).

### Histology

For EdU staining tissues were embedded in paraffin and permeabilized with 0.5 % Triton X-100 (Sigma-Aldrich) in PBS after dewaxing. Slides were washed with 3% BSA-PBS solution for 7 min. Click-iT™ Plus reaction cocktail was mixed according to the manufacturer’s instructions, with an antibody concentration of 1:20. Cells were incubated twice for 30 min at RT. After washing in 3% BSA-PBS, nuclei were stained with SYBR Green (Invitrogen).

For cytochrome c oxidase staining frozen tissue sections were thawed at RT for 20 min. To minimize time delays and reduce staining variability, only 1 slide per group and 1 control were stained per round. Per sample, a total of 3 slides were stained and analyzed. Sections were incubated uncovered with COX media (1X 3,3’-diaminobenzidine, 1mg/ml bovine cytochrome c, 2 µg/ml bovine catalase (Sigma-Aldrich)) at 37°C for 40 min in a humidity chamber. For the control slides, enzyme activity was blocked with 1 mg/ml sodium azide (Sigma-Aldrich). Slides were washed four times with 10X PBS, dehydrated with decreasing concentrations of ethanol and xylol and mounted with entellan (Sigma-Aldrich). Slides were scanned with a 3DHistech Pannoramic 250 Flash II.

### MitoSpy™ flow cytometry assay

Prior to adding fixable viability dye (eBioscience), cells were incubated for 35 min at 37°C in 200 µl MitoSpy™ orange CMTMRos (1:2000 in incomplete IMDM; Biolegend). Subsequently, cells were washed twice with PBS and stained with fixable viability dye (eBioscience) for 20 min and extracellular Epcam marker (Biolegend) for 15 min on ice. After washing with FACS buffer (2% FCS, 2 mM EDTA), cells were fixed and permeabilized with the Foxp3/Transcription Factor Staining Buffer Set (eBioscience) following the manufacturer’s protocol.

### Cytochrome c oxidase staining

Tissues were embedded in OCT embedding matrix (CellPath) and cut in 7 µm sections on a Cryostat Microm HM 560 at −20°C, dried for 10 min at RT and frozen at −20°C without fixation. For cytochrome c oxidase staining sections were thawed at RT for 20 min. Sections were incubated uncovered with COX media (1X 3,3’-diaminobenzidine, 1 mg/ml bovine cytochrome c, 2 µg/ml bovine catalase (Sigma-Aldrich)) at 37°C for 40 min in a humidity chamber. For the control slides, enzyme activity was blocked with 1 mg/ml sodium azide (Sigma-Aldrich). Slides were washed four times with 10X PBS, dehydrated with decreasing concentrations of ethanol and xylol and mounted with entellan (Sigma-Aldrich).

### Intestinal organoid culture

The terminal ileum was opened longitudinally and washed in PSG (1:100 penicillin/streptomycin 10’000 units (Gibco) and 70 µg/ml gentamicin (Gibco) in 1X PBS) to remove intestinal contents. The tissue was cut into 2 cm pieces and washed in PSG 10 times. To dissociate villi and crypts from the lamina propria, pieces were incubated on a horizontal shaker for 15 min at RT in 15 ml 1X PBS (Gibco) supplemented with 7.5 mM EDTA and 10 µM Y27632 (ROCK inhibitor; Sigma-Aldrich) and 10 mM HEPES (Gibco). Villi were subsequently discarded by vigorous vortexing over three consecutive rounds with fresh 1X PBS. The resulting crypt fractions were collected in complete DMEM/F12^+^ Glutamax media (supplemented with 10% FCS, 10 mM HEPES) by passing through a 100 µm cell strainer (VWR) and 70 µm cell strainer (VWR). Crypts were counted and resuspended in 1 part matrigel (Corning) and 1 part WENR media (1:100 N2 supplement (Gibco), 1:50 B27 supplement (Gibco), 0.1 mM HEPES (Gibco), 10 units penicillin/streptomycin (Gibco), 240 µg/ml gentamicin (Gibco), 0.1 µg/ml EGF, 0.1 µg/ml recombinant murine noggin (Peprotech), 0.5 µg/ml recombinant murine R-Spondin (R&D systems), 10 µg/ml Y27632 (Sigma-Aldrich), 0.25 µg/ml recombinant murine 3a (Peprotech), 10 % FCS). Approximately 500 crypts were seeded per 50 µl matrigel droplet in 24 well tissue culture plates (TPP) for all other analyses. After the matrigel had solidified, WENR media was added to each culture well. Media was changed every second day and ENR media was added when organoids were ready to differentiate. ENR media had the same compositions as WENR media but lacked recombinant murine Wnt3a and Y27632. For treatment 10 mM palmitoleic acid (Sigma-Aldrich) was supplemented in ENR media for 3 h. Organoids were then harvested with ice cold media to dissolve the matrigel. They were collected in Trizol after centrifugation (500 g, 5 min, 4°C) and snap-frozen.

### Metabolomic analysis

Intestinal contents were collected and snap-frozen in liquid nitrogen. Small intestinal epithelial cells were isolated as described above and snap-frozen as pellet. Metabolites were extracted with 20 parts of 80°C Millipore water per sample weight. Samples were shortly put on a thermomixer at 80°C before homogenization for 2 min at 30 Hz (Retsch MM 400). Extraction was performed on the thermomixer at 80°C for 3 min at maximal speed and with vortexing for 10 seconds every minute. After centrifugation (3 min, 20’000 g, RT) the supernatant was collected, diluted and sealed in 96-well plates.

Tissue samples were diluted 1/10 and gut content samples had a final dilution of 1/100 in MS grade ddH2O prior to injection. An Agilent 6550 QTOF mass spectrometer (Agilent) was used with the method described in Fuhrer et al. ^70^. Samples were run in two batches and injected in a randomized sequence using a Gerstel MPS 3 autosampler (Gerstel). The instrument was run at 4 GHz in negative ionization MS1 mode collecting high resolution mass spectra between 50 and 1000 mass/charge (m/z) ratio. The mobile phase was 60:40 isopropanol:water adjusted to pH 9 using 4 mM NH4F and supplemented with 10 nM hexakis (1H, 1H, 3H-tetrafluoropropoxy)phosphazine and 80 nM taurochloric acid for online mass calibration. Spectra from the two batches (tissue and content) sample runs were aligned and centroided using superalign in matlab. Ions were annotated by matching their measured mass with reference compounds derived from the Human Metabolome Database (HMDB 4.0) subselected for endogenous compounds, fecal and bacterial metabolites and allowing a tolerance of 1 mDa. The resulting data matrix contained 1’508 ions that could be matched to the deprotonated metabolites from the database. All m/z peaks that remained unmatched were discarded. Ions were further TIC-normalized and the further data processing was performed in a customized R pipeline (https://github.com/DorotheeBerthold/maternal-gut-metabolites). The obtained metabolite identifiers were converted to SMILES and submitted to Classyfire ^71^ to retrieve metabolite classes. In case multiple classes were annotated for a single feature, a ranking system came into place, selecting the most likely class based on co-occurrence among other features.

### Data availability

ATAC sequencing, RNA sequencing and whole genome bisulfite sequencing (WGBS) data will be made available at the time of publication on the NCBI Sequencing Read Archive under the following BioProject numbers: ATAC-Seq – PRJNA1507342; RNA-Seq – PRJNA1507620; WGBS – PRJNA1509478.

Raw mass spectrometry data is deposited on massive.ucsd.edu under MSV000100948.

### Quantification and statistical analysis

If not stated otherwise, an unpaired parametric Student’s t-test was performed to compare two experimental groups. These tests were performed in GraphPad Software v.9.1.2 or in R v.4.1.2. If p-values are indicated with asterisks, the following applies * p < 0.05, ** p ≤ 0.01, *** p ≤ 0.001, **** p ≤ 0.0001. Error bars indicate mean ± standard deviation. Figures were created with R v.4.1.2 and subsequently merged in Adobe Illustrator v.24.2.3.

## RESOURCE AVAILABILITY

### Lead contact

Requests for further information and resources should be directed to and will be fulfilled by the lead contact, Stephanie Ganal-Vonarburg.

### Materials availability

This study did not generate new unique reagents. *E. coli* HA107 used in this study will be made available on request by Andrew Macpherson or the lead contact, but we may require a completed materials transfer agreement.

### Data and code availability

RNA-seq, whole-genome bisulfite sequencing and ATAC-seq raw data have been deposited at on the NCBI Sequencing Read Archive under the following BioProjects: PRJNA1507342, PRJNA1507620, PRJNA1509478, and are publicly available as of the date of publication. Raw mass spectrometry data have been deposited on massive.ucsd.edu under MSV000100948, and are publicly available as of the date of publication.

All original code and any additional information required to reanalyze the data reported in this paper is available from the lead contact upon request.

## ACKNOWLEDGMENTS

We are grateful for support by Pamela Nicholson and the team of the Next Generation Sequencing Facility of the University of Bern and by the Clean Mouse Facility, EAC, which is supported by the Genaxen Foundation, Inselspital and the University of Bern. We acknowledge Adriano Rutz for his assistance with the annotation of metabolite classes and Steven Misztal for guidance in the in silico flow cytometry.

## Funding

Swiss National Science Foundation grant 310030_212511 (SCGV)

Foundation for Experimental Biomedicine Peter Hans Hofschneider Stiftungsprofessur (SCGV)

Swiss National Foundation Sinergia grant CRSII5_205965 (US, AJM)

Swiss National Science Foundation MD-PhD grant 323530_199385 (SC)

European Research Council grant H2020 ERC-2016-ADG HHMM_Neonates Grant agreement: 742195 (AJM)

## AUTHOR CONTRIBUTIONS

Conceptualization: SCGV, CK, AJM

Methodology: SCGV, CK, ST, AJM, SC, IK, RB, HL, SM, ST, US, JCF

Investigation: CK, SC, ST, NFT, IK, HL, DLB, AW, MGA, SCGV, JCF

Visualization: CK, SGV, IK, DLB, JCF

Funding acquisition: SCGV, AJM

Project administration: SCGV

Supervision: SCGV

Writing – original draft: SCGV, CK, AJM

Writing – review & editing: SCGV, CK, AJM, DLB, SC, ST, JCF

## DECLARATION OF INTERESTS

The authors declare that they have no competing interests.

## DECLARATION OF GENERATIVE AI AND AI-ASSISTED TECHNOLOGIES IN THE WRITING PROCESS

During the preparation of this work, the authors used ChatGPT (OpenAI) in order to improve English language style. After using this tool or service, the authors reviewed and edited the content as needed and take full responsibility for the content of the publication.

## SUPPLEMENTAL INFORMATION

**Data S1. (separate file):** List of genes in subclusters shown in Figure S5B.

**Data S2. (separate file):** Metabolite annotations in clusters shown in Figure 5.

---

**Figure S1.**
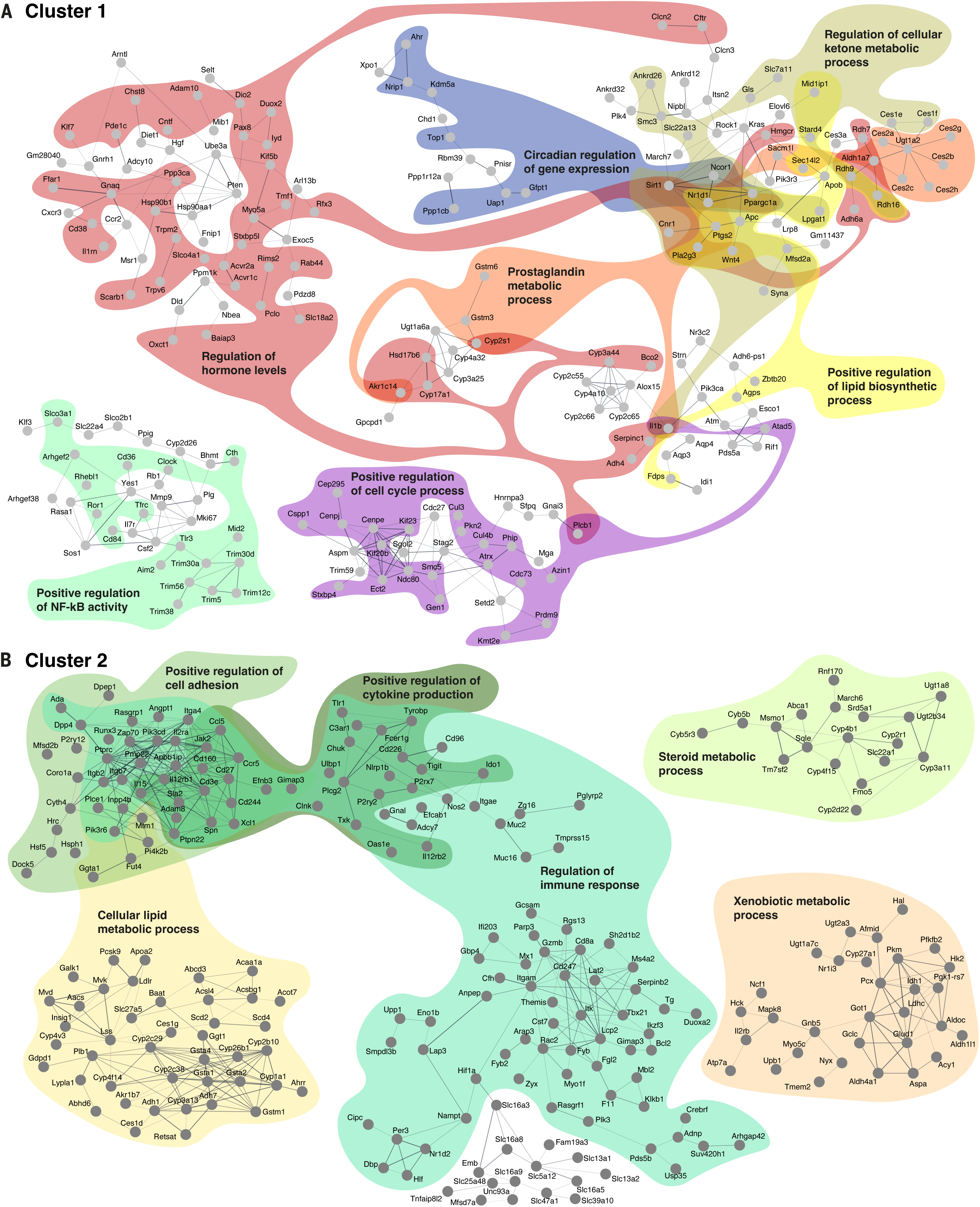
Genes of cluster 1 and 2 selected and clustered according to GO analysis. Differentially expressed genes corresponding to the clusters in Figure 1B were subjected to GO analysis and subsequently illustrated according to STRING pathways. **A.** Genes in cluster 1 and genes in **B.** cluster 2 are shown.

**Figure S2.**
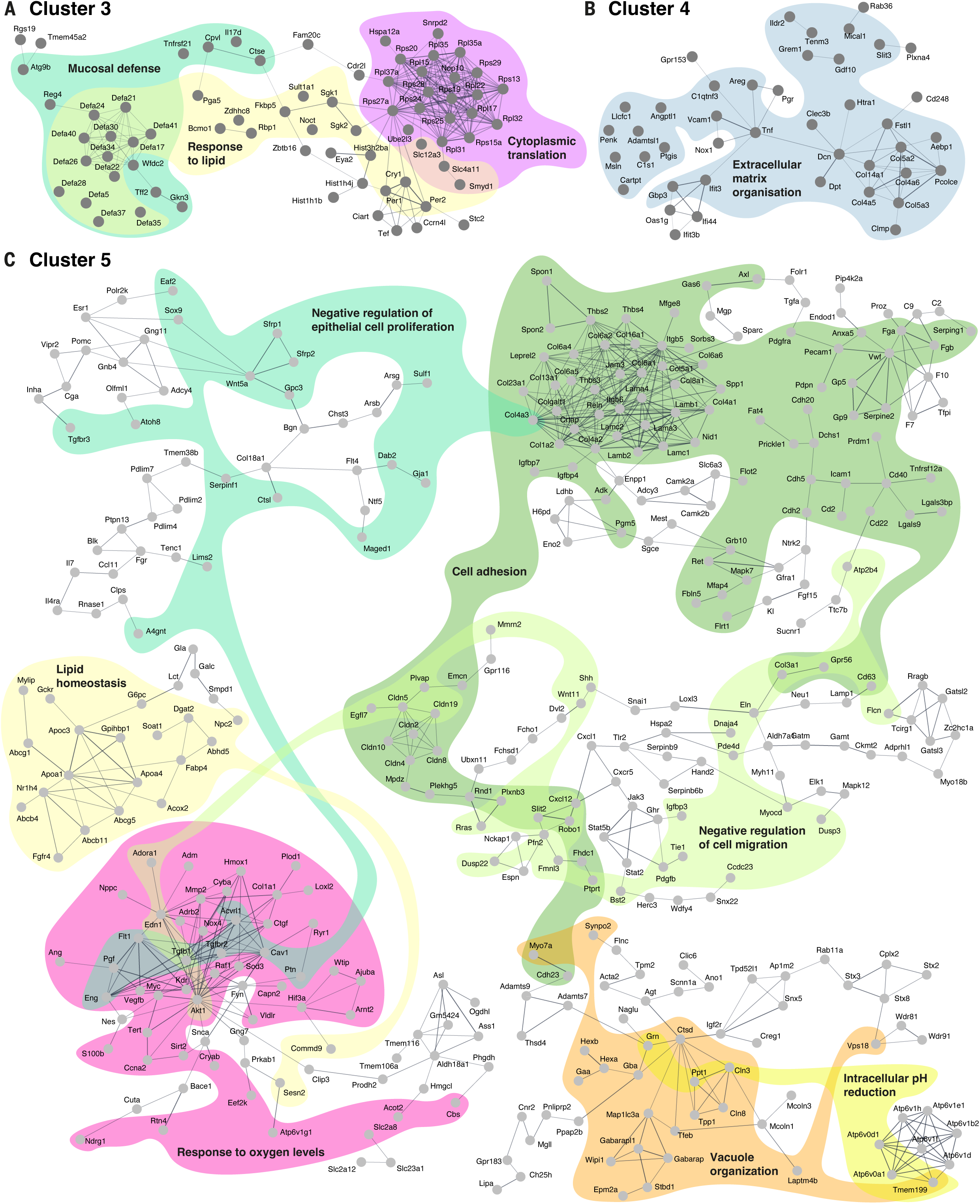
Genes of cluster 3, 4, and 5 selected and clustered according to GO analysis. Differentially expressed genes corresponding to the clusters in Figure 1B were subjected to GO analysis and subsequently illustrated according to STRING pathways. **A.** Genes in cluster 3, in **B.** cluster 4 and in **C.** cluster 5 are displayed.

**Figure S3.**
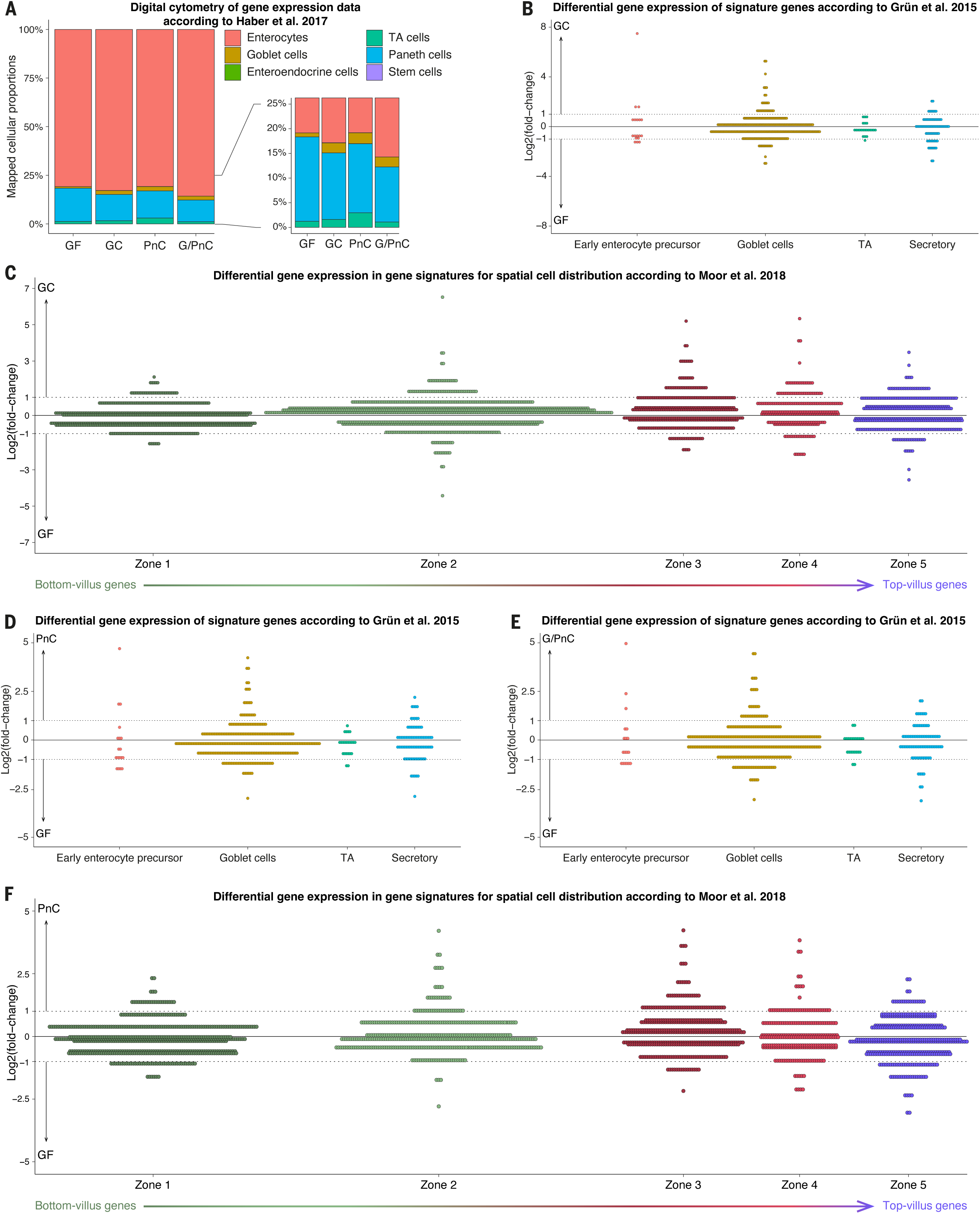
In silico estimation of cellular distribution in small intestines of germ-free, gestationally colonized and postnatally colonized mice. **A.** Bar plot displaying cell distribution computed with the online tool CIBERSORTx ^58^ by imputing the RNA sequencing data. Data was mapped on published single-cell data of murine small intestinal epithelial cells ^59^. **B.** Differential gene expression (log2-fold-change values) between germ-free (GF) and gestationally colonized (GC) offspring was mapped for signature genes within the early enterocyte precursors, goblet cells, transit amplifying cells (TA) and secretory lineage according to ^72^. No cut-off for the p-value was used. **C.** Differential gene expression (log2-fold-change values) between GF and GC offspring mapped on zonation signature genes according to ^73^. No cut-off for the p-value was used. **D.** Differential gene expression (log2-fold-change values) between GF compared to PnC mice. No cut-off for the p-value was used. **E.** Differential gene expression (log2-fold-change values) between GF and G/PnC mice were mapped on signature genes of early enterocyte precursors, goblet cells, transit amplifying cells (TA) and secretory lineages according to ^72^. No cut-off for the p-value was used. **F.** Differential gene expression (log2-fold-change values) between GF and PnC mice mapped on zonation signature genes according to ^73^. No cut-off for the p-value was used.

**Figure S4.**
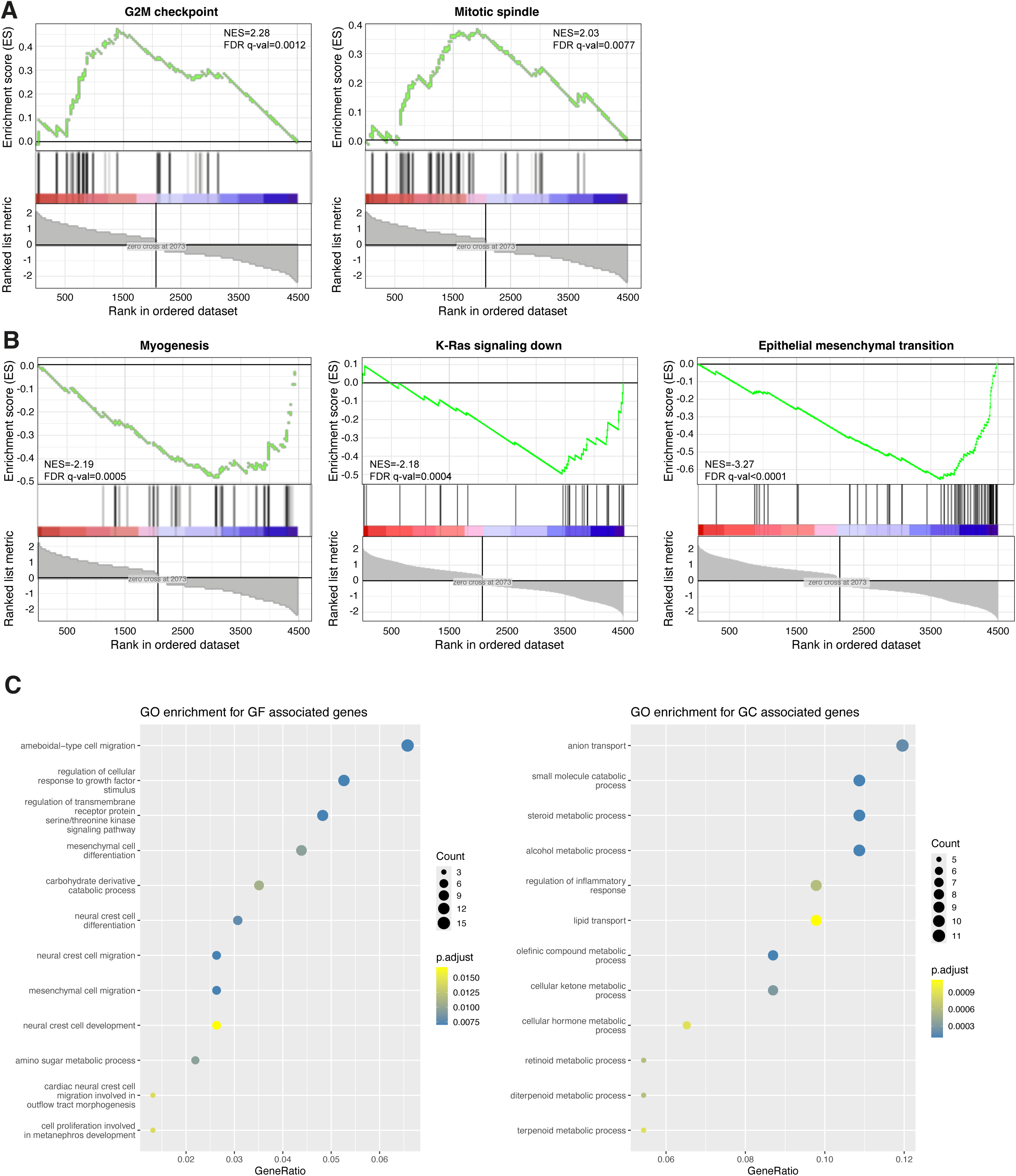
Gene set enrichment analysis of differentially expressed genes between germ-free and gestationally colonized mice. **A, B.** Gene sets significantly (FDR < 0.05) enriched in GC (A) offspring or in GF (B) offspring. Enrichment scores (ES) are displayed on the y-axis. Ranked genes within the gene sets are visualized as bars on the x-axis. The green line demonstrates the correlation the ranked genes and the enrichment score, with positive values are sets with higher enrichment score in the GC group, while negative values represent sets enriched in GF mice. The degree of correlation is also demonstrated with the colored bar (red for positive and blue for negative correlation). **C.** GO enrichment analysis on genes showing overlap in differential gene expression and ATAC differentially accessible peaks in P21 small IECs in GF versus GC offspring. GF associated genes: gene expression increased (adj p-value < 0.05 and fold-change > 2) and enriched ATAC peak (adj p-value < 0.05 and fold-change > 0) in GF compared to GC group; GC associated genes: gene expression increased (adj p-value < 0.05 and fold-change < −2) and enriched ATAC peak (adj p-value < 0.05 and fold-change < 0) in GC compared to GF groups (see overlap in Figure 2D).

**Figure S5.**
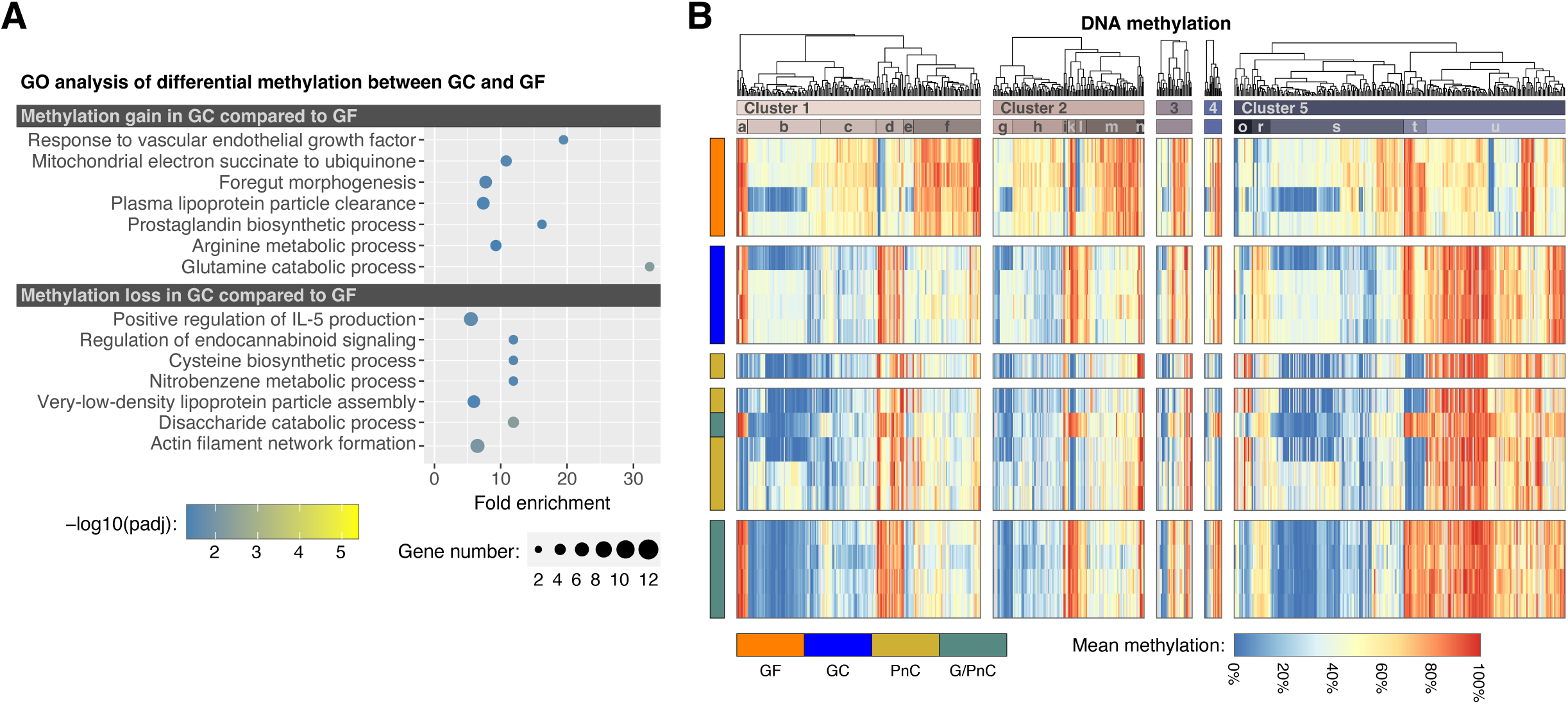
DNA methylation analysis on P21 - GO analysis and overlap with RNA-seq results. **A.** GO analysis of the genes with a methylation gain or loss (≥ 25%; padj < 0.05) in GC compared to GF. Nodes are colored according to the padj and the size is relative to the number of genes per GO term. **B.** Mean DNA methylation of differentially methylated genes (≥ 25%; padj < 0.05) represented as individual heat maps per gene clusters from panel in Figure 1B. For the mean methylation level per gene, CpGs were restricted to those within +/- 20 kb from the TSS with intergenic CpGs excluded. Remaining CpGs were then averaged by gene.

**Figure S6.**
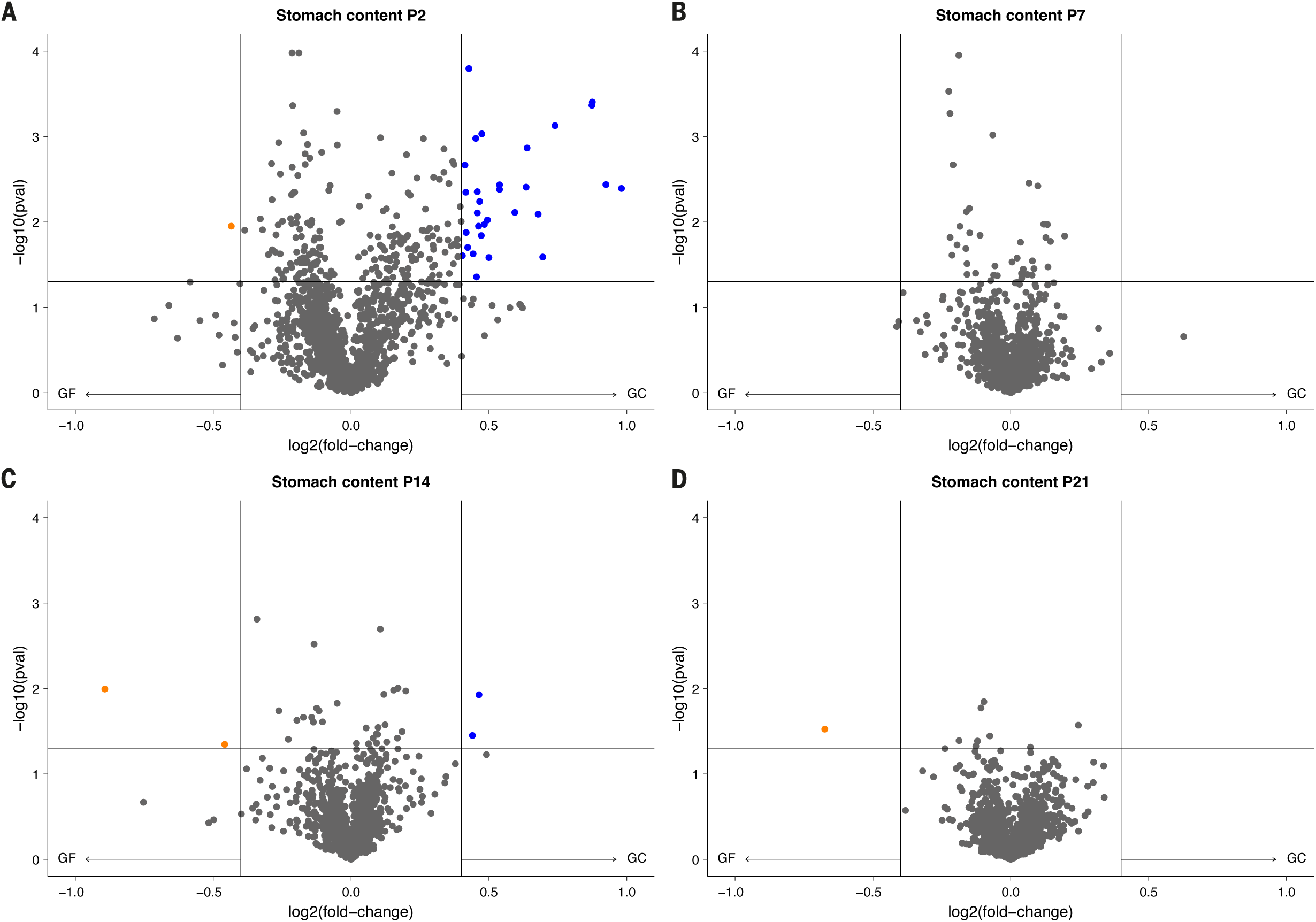
Time trajectory of differential metabolic content in the stomach of germ-free and gestationally colonized offspring early in life. Volcano plot demonstrating differences in the metabolome between germ-free (GF) and gestationally colonized (GC) offspring (fold-change ≥ 0.4; p-value < 0.05) on **A.** postnatal day 2 (P2), **B.** postnatal day 7 (P7), **C.** postnatal day 14 (P14) and **D.** postnatal day 21 (P21).

**Figure S7.**
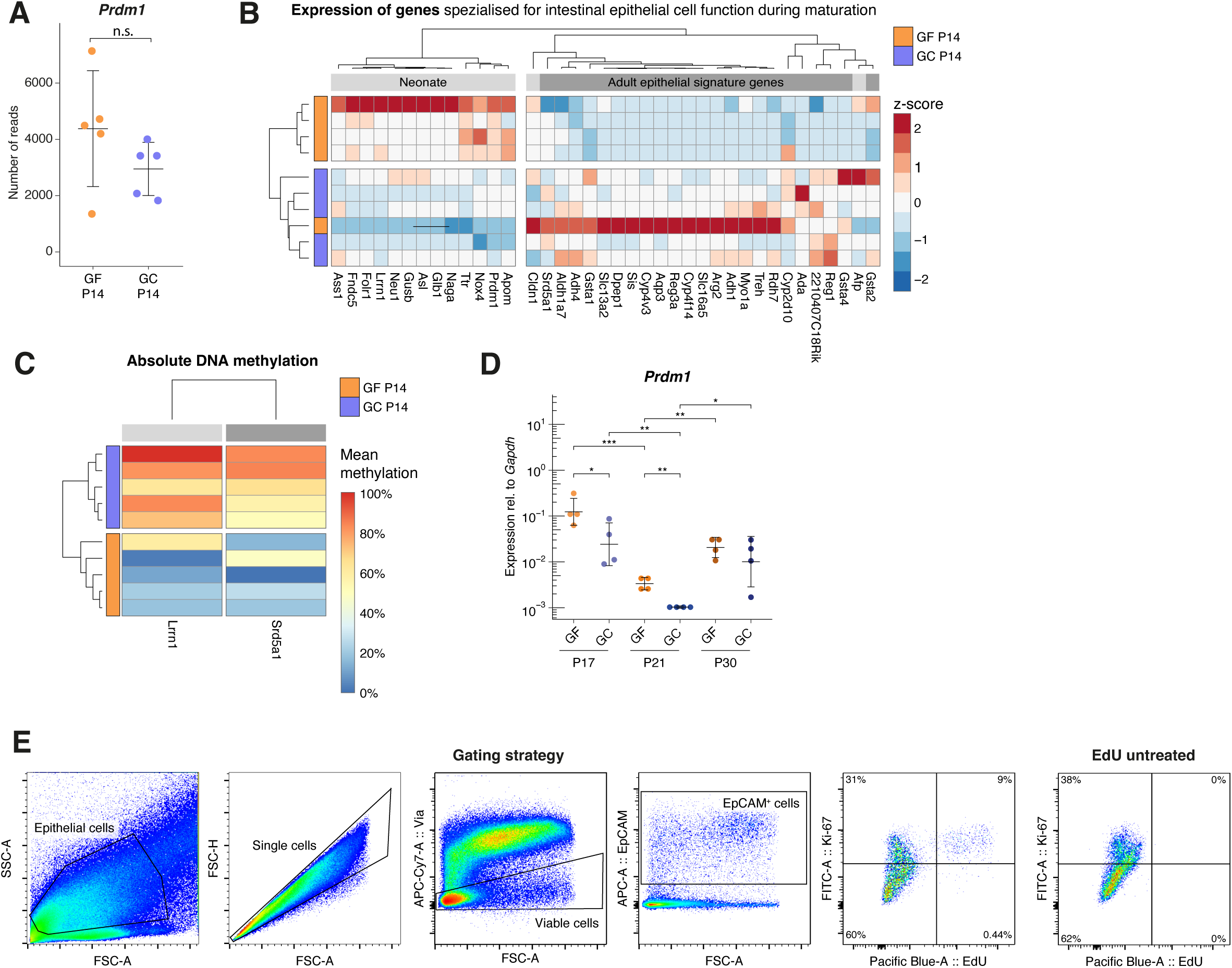
Effect of the maternal microbiota on BLIMP-1 regulated epithelial transition in the 14-day-old offspring. **A.** Numbers of normalized reads of the gene encoding for BLIMP-1 (*Prdm1*) in 14-day-old pups born to germ-free or gestationally colonized dams. **B.** Heat map illustrating the relative expression profile of genes responding to BLIMP-1 according to ^11,12^) and ^11,12^) in GF and GC P14 neonates. **C.** The absolute DNA methylation of BLIMP-1 regulated genes, which are differentially methylated (≥ 25%; padj < 0.05) between 14-day-old pups. D. RT-qPCR results for *Prdm1* expression in small intestinal epithelial cells isolated from GF and GF offspring at postnatal days 17, 21 and 30. **E.** Flow cytometry strategy to gate for EpCAM^+^EdU^+^ and EpCAM^+^Ki-67^+^ cells in the small intestine after i.p. administration of EdU and representative final gate for a sample originating from an EdU untreated control mouse.

**Figure S8.**
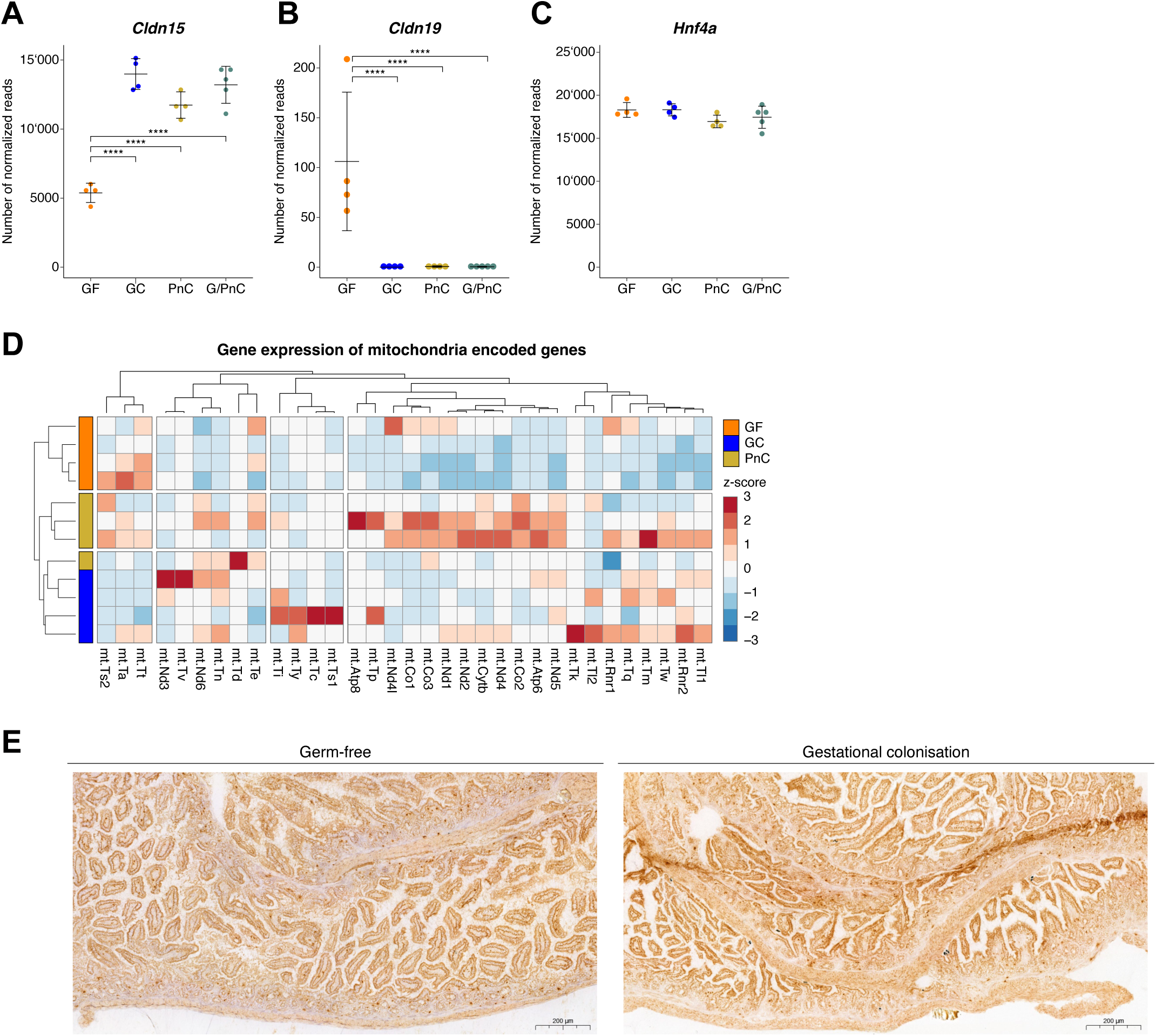
Gene expression associated to the epithelial barrier in 21-day-old pups. **A-C.** Numbers of normalized reads in the RNA sequencing analysis of gene encoding for the barrier forming claudin 15 (A), the pore-forming claudin 19 (B) and the regulatory transcription factor Hnf4a (C) in 21-day-old germ-free pups (GF), pups born to gestationally colonized dams (GC) and pups treated with HA107 during weaning either born to a germ-free dam (PnC) or to a gestationally colonized dam (GC). The error bars display the mean ± standard deviation. Asterisks indicate q-values with **** q ≤ 0.0001. **D.** Heat map with differential gene expression of genes on the mitochondrial genome. The color scale indicates the relative expression profiles per sample. **E.** Representative cryosections of the small intestine after enzymatic staining for cytochrome c oxidase activity in P21 old GF and GC offspring.

**Figure S9.**
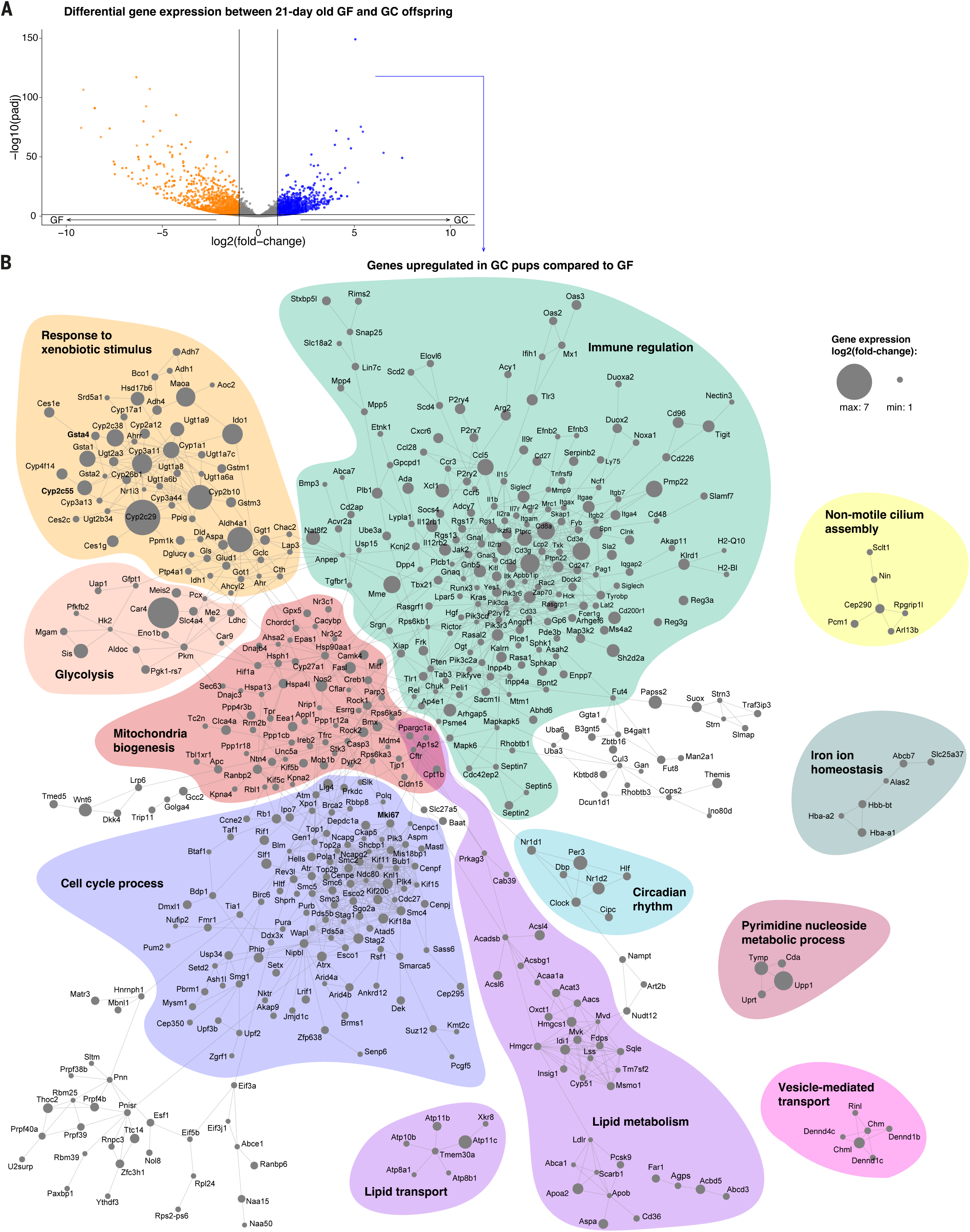
Differential gene expression of 21-day-old germ-free and gestationally primed offspring. **A.** Volcano plot visualizing transcriptomic differences between 21-day-old mice born to gestationally colonized (GC) or germ-free (GF) dams (fold-change ≥ 2; padj < 0.05). **B.** STRING enrichment analysis of upregulated genes in small intestinal epithelial cells upon gestational colonization (fold-change ≥ 2; padj < 0.05). The size of the nodes displays the log2(fold-change) in the transcriptomic data (confidence cutoff = 0.9).

## Notes

### Competing Interest Statement

The authors have declared no competing interest.

## REFERENCES

1. Torow, N., and Hornef, M.W. (2017). The Neonatal Window of Opportunity: Setting the Stage for Life-Long Host-Microbial Interaction and Immune Homeostasis. J Immunol 198, 557–563. 10.4049/jimmunol.1601253.

2. Hornef, M.W., and Torow, N. (2020). ’Layered immunity’ and the ‘neonatal window of opportunity’-timed succession of non-redundant phases to establish mucosal host-microbial homeostasis after birth. Immunology 159, 15–25. 10.1111/imm.13149.

3. Walter, J., and Hornef, M.W. (2021). A philosophical perspective on the prenatal in utero microbiome debate. Microbiome 9, 5. 10.1186/s40168-020-00979-7.

4. Gomez de Agüero, M., Ganal-Vonarburg, S.C., Fuhrer, T., Rupp, S., Uchimura, Y., Li, H., Steinert, A., Heikenwalder, M., Hapfelmeier, S., Sauer, U., et al. (2016). The maternal microbiota drives early postnatal innate immune development. Science 351, 1296–1302. 10.1126/science.aad2571.

5. Lim, A.I., McFadden, T., Link, V.M., Han, S.J., Karlsson, R.M., Stacy, A., Farley, T.K., Lima-Junior, D.S., Harrison, O.J., Desai, J.V., et al. (2021). Prenatal maternal infection promotes tissue-specific immunity and inflammation in offspring. Science 373. 10.1126/science.abf3002.

6. Kimura, I., Miyamoto, J., Ohue-Kitano, R., Watanabe, K., Yamada, T., Onuki, M., Aoki, R., Isobe, Y., Kashihara, D., Inoue, D., et al. (2020). Maternal gut microbiota in pregnancy influences offspring metabolic phenotype in mice. Science 367, eaaw8429. 10.1126/science.aaw8429.

7. Uchimura, Y., Fuhrer, T., Li, H., Lawson, M.A., Zimmermann, M., Yilmaz, B., Zindel, J., Ronchi, F., Sorribas, M., Hapfelmeier, S., et al. (2018). Antibodies Set Boundaries Limiting Microbial Metabolite Penetration and the Resultant Mammalian Host Response. Immunity 49, 545–559.e545. 10.1016/j.immuni.2018.08.004.

8. Tsang, D.K.L., Wang, R.J., De Sa, O., Ayyaz, A., Foerster, E.G., Bayer, G., Goyal, S., Trcka, D., Ghoshal, B., Wrana, J.L., et al. (2022). A single cell survey of the microbial impacts on the mouse small intestinal epithelium. Gut Microbes 14, 2108281. 10.1080/19490976.2022.2108281.

9. Camp, J.G., Frank, C.L., Lickwar, C.R., Guturu, H., Rube, T., Wenger, A.M., Chen, J., Bejerano, G., Crawford, G.E., and Rawls, J.F. (2014). Microbiota modulate transcription in the intestinal epithelium without remodeling the accessible chromatin landscape. Genome Res 24, 1504–1516. 10.1101/gr.165845.113.

10. Qin, Y., Roberts, J.D., Grimm, S.A., Lih, F.B., Deterding, L.J., Li, R., Chrysovergis, K., and Wade, P.A. (2018). An obesity-associated gut microbiome reprograms the intestinal epigenome and leads to altered colonic gene expression. Genome Biol 19, 7. 10.1186/s13059-018-1389-1.

11. Muncan, V., Heijmans, J., Krasinski, S.D., Büller, N.V., Wildenberg, M.E., Meisner, S., Radonjic, M., Stapleton, K.A., Lamers, W.H., Biemond, I., et al. (2011). Blimp1 regulates the transition of neonatal to adult intestinal epithelium. Nature Communications 2, 452. 10.1038/ncomms1463.

12. Harper, J., Mould, A., Andrews, R.M., Bikoff, E.K., and Robertson, E.J. (2011). The transcriptional repressor Blimp1/Prdm1 regulates postnatal reprogramming of intestinal enterocytes. Proc Natl Acad Sci U S A 108, 10585–10590. 10.1073/pnas.1105852108.

13. Zhang, K., Dupont, A., Torow, N., Gohde, F., Leschner, S., Lienenklaus, S., Weiss, S., Brinkmann, M.M., Kuhnel, M., Hensel, M., et al. (2014). Age-dependent enterocyte invasion and microcolony formation by Salmonella. PLoS Pathog 10, e1004385. 10.1371/journal.ppat.1004385.

14. Holmes, J.L., Van Itallie, C.M., Rasmussen, J.E., and Anderson, J.M. (2006). Claudin profiling in the mouse during postnatal intestinal development and along the gastrointestinal tract reveals complex expression patterns. Gene Expression Patterns 6, 581–588. 10.1016/j.modgep.2005.12.001.

15. Gleeson, J.P., Fein, K.C., Chaudhary, N., Doerfler, R., Newby, A.N., and Whitehead, K.A. (2021). The enhanced intestinal permeability of infant mice enables oral protein and macromolecular absorption without delivery technology. International Journal of Pharmaceutics 593, 120120. 10.1016/j.ijpharm.2020.120120.

16. Patel, R.M., Myers, L.S., Kurundkar, A.R., Maheshwari, A., Nusrat, A., and Lin, P.W. (2012). Probiotic Bacteria Induce Maturation of Intestinal Claudin 3 Expression and Barrier Function. The American Journal of Pathology 180, 626–635. 10.1016/j.ajpath.2011.10.025.

17. Remis, N.N., Wiwatpanit, T., Castiglioni, A.J., Flores, E.N., Cantu, J.A., and Garcia-Anoveros, J. (2014). Mucolipin co-deficiency causes accelerated endolysosomal vacuolation of enterocytes and failure-to-thrive from birth to weaning. PLoS Genet 10, e1004833. 10.1371/journal.pgen.1004833.

18. Moretti, D.B., Nordi, W.M., Lima, A.L., Pauletti, P., and Machado-Neto, R. (2013). Enterocyte IgG uptake in the small intestine of goat kids during the period of passive immunity acquisition. Small Ruminant Research 114, 182–187. 10.1016/j.smallrumres.2013.05.012.

19. Moxey, P.C., and Trier, J.S. (1979). Development of villus absorptive cells in the human fetal small intestine: A morphological and morphometric study. The Anatomical Record 195, 463–482. 10.1002/ar.1091950307.

20. Price, A.E., Shamardani, K., Lugo, K.A., Deguine, J., Roberts, A.W., Lee, B.L., and Barton, G.M. (2018). A Map of Toll-like Receptor Expression in the Intestinal Epithelium Reveals Distinct Spatial, Cell Type-Specific, and Temporal Patterns. Immunity 49, 560–575 e566. 10.1016/j.immuni.2018.07.016.

21. Fulde, M., Sommer, F., Chassaing, B., van Vorst, K., Dupont, A., Hensel, M., Basic, M., Klopfleisch, R., Rosenstiel, P., Bleich, A., et al. (2018). Neonatal selection by Toll-like receptor 5 influences long-term gut microbiota composition. Nature 560, 489–493. 10.1038/s41586-018-0395-5.

22. Pott, J., Stockinger, S., Torow, N., Smoczek, A., Lindner, C., McInerney, G., Backhed, F., Baumann, U., Pabst, O., Bleich, A., and Hornef, M.W. (2012). Age-dependent TLR3 expression of the intestinal epithelium contributes to rotavirus susceptibility. PLoS Pathog 8, e1002670. 10.1371/journal.ppat.1002670.

23. Yu, D.-H., Gadkari, M., Zhou, Q., Yu, S., Gao, N., Guan, Y., Schady, D., Roshan, T.N., Chen, M.-H., Laritsky, E., et al. (2015). Postnatal epigenetic regulation of intestinal stem cells requires DNA methylation and is guided by the microbiome. Genome Biology 16, 211. 10.1186/s13059-015-0763-5.

24. Kaaij, L.T.J., van de Wetering, M., Fang, F., Decato, B., Molaro, A., van de Werken, H.J.G., van Es, J.H., Schuijers, J., de Wit, E., de Laat, W., et al. (2013). DNA methylation dynamics during intestinal stem cell differentiation reveals enhancers driving gene expression in the villus. Genome Biology 14, R50. 10.1186/gb-2013-14-5-r50.

25. Pan, W.-H., Sommer, F., Falk-Paulsen, M., Ulas, T., Best, P., Fazio, A., Kachroo, P., Luzius, A., Jentzsch, M., Rehman, A., et al. (2018). Exposure to the gut microbiota drives distinct methylome and transcriptome changes in intestinal epithelial cells during postnatal development. Genome Medicine 10, 27. 10.1186/s13073-018-0534-5.

26. Yang, L., Peery, R.C., Zhou, S., Chen, X., Farmer, L.M., Gutierrez, F., Fowler, S., Zhang, L., Salamat, J.M., Riggins, K., et al. (2026). Weaning drives microbiome-mediated epigenetic regulation to shape immune memory in mice. Nat Microbiol 11, 1064–1079. 10.1038/s41564-026-02295-6.

27. Hapfelmeier, S., Lawson, M.A.E., Slack, E., Kirundi, J.K., Stoel, M., Heikenwalder, M., Cahenzli, J., Velykoredko, Y., Balmer, M.L., Endt, K., et al. (2010). Reversible Microbial Colonization of Germ-Free Mice Reveals the Dynamics of IgA Immune Responses. Science 328, 1705–1709. doi:10.1126/science.1188454.

28. Ternet, C., and Kiel, C. (2021). Signaling pathways in intestinal homeostasis and colorectal cancer: KRAS at centre stage. Cell Communication and Signaling 19, 31. 10.1186/s12964-021-00712-3.

29. Kalluri, R., and Weinberg, R.A. (2009). The basics of epithelial-mesenchymal transition. J Clin Invest 119, 1420–1428. 10.1172/JCI39104.

30. Venkatraman, A., Yu, W., Nitkin, C., and Sampath, V. (2021). Intestinal Stem Cell Development in the Neonatal Gut: Pathways Regulating Development and Relevance to Necrotizing Enterocolitis. Cells 10. 10.3390/cells10020312.

31. Gehart, H., and Clevers, H. (2019). Tales from the crypt: new insights into intestinal stem cells. Nat Rev Gastroenterol Hepatol 16, 19–34. 10.1038/s41575-018-0081-y.

32. Guerin, L.N., Scott, T.J., Yap, J.A., Johansson, A., Puddu, F., Charlesworth, T., Yang, Y., Simmons, A.J., Lau, K.S., Ihrie, R.A., and Hodges, E. (2025). Temporally discordant chromatin accessibility and DNA demethylation define short- and long-term enhancer regulation during cell fate specification. Cell Rep 44, 115680. 10.1016/j.celrep.2025.115680.

33. de Santa Barbara, P., van den Brink, G.R., and Roberts, D.J. (2003). Development and differentiation of the intestinal epithelium. Cellular and Molecular Life Sciences CMLS 60, 1322–1332. 10.1007/s00018-003-2289-3.

34. Khan, N., and Asif, A.R. (2015). Transcriptional regulators of claudins in epithelial tight junctions. Mediators Inflamm 2015, 219843. 10.1155/2015/219843.

35. Ikegami, S., Maeda, K., Urano, T., Mu, J., Nakamura, M., Yamamura, T., Sawada, T., Ishikawa, E., Yamamoto, K., Muto, H., et al. (2024). Monoclonal Antibody Against Mature Interleukin-18 Ameliorates Colitis in Mice and Improves Epithelial Barrier Function. Inflammatory Bowel Diseases 30, 1353–1366. 10.1093/ibd/izad292.

36. Kotler, B.M., Kerstetter, J.E., and Insogna, K.L. (2013). Claudins, dietary milk proteins, and intestinal barrier regulation. Nutr Rev 71, 60–65. 10.1111/j.1753-4887.2012.00549.x.

37. Sugimoto, K., and Chiba, H. (2021). The claudin–transcription factor signaling pathway. Tissue Barriers 9, 1908109. 10.1080/21688370.2021.1908109.

38. Luettig, J., Rosenthal, R., Barmeyer, C., and Schulzke, J.D. (2015). Claudin-2 as a mediator of leaky gut barrier during intestinal inflammation. Tissue Barriers 3, e977176. 10.4161/21688370.2014.977176.

39. Popov, L.-D. (2020). Mitochondrial biogenesis: An update. Journal of Cellular and Molecular Medicine 24, 4892–4899. 10.1111/jcmm.15194.

40. Villena, J.A. (2015). New insights into PGC-1 coactivators: redefining their role in the regulation of mitochondrial function and beyond. The FEBS Journal 282, 647–672. 10.1111/febs.13175.

41. Berger, E., Rath, E., Yuan, D., Waldschmitt, N., Khaloian, S., Allgäuer, M., Staszewski, O., Lobner, E.M., Schöttl, T., Giesbertz, P., et al. (2016). Mitochondrial function controls intestinal epithelial stemness and proliferation. Nature Communications 7, 13171. 10.1038/ncomms13171.

42. Whitt, J., Woo, V., Lee, P., Moncivaiz, J., Haberman, Y., Denson, L., Tso, P., and Alenghat, T. (2018). Disruption of Epithelial HDAC3 in Intestine Prevents Diet-Induced Obesity in Mice. Gastroenterology 155, 501–513. 10.1053/j.gastro.2018.04.017.

43. Fahur Bottino, G., Bonham, K.S., Patel, F., McCann, S., Zieff, M., Naspolini, N., Ho, D., Portlock, T., Joos, R., Midani, F.S., et al. (2025). Early life microbial succession in the gut follows common patterns in humans across the globe. Nat Commun 16, 660. 10.1038/s41467-025-56072-w.

44. D’Aquila, P., Carelli, L.L., De Rango, F., Passarino, G., and Bellizzi, D. (2020). Gut Microbiota as Important Mediator Between Diet and DNA Methylation and Histone Modifications in the Host. Nutrients 12. 10.3390/nu12030597.

45. Ouzounis, C.A., and Karp, P.D. (2000). Global properties of the metabolic map of Escherichia coli. Genome Res 10, 568–576. 10.1101/gr.10.4.568.

46. Stevens, J.F., and Maier, C.S. (2016). The Chemistry of Gut Microbial Metabolism of Polyphenols. Phytochem Rev 15, 425–444. 10.1007/s11101-016-9459-z.

47. Zheng, L., Kelly, C.J., Battista, K.D., Schaefer, R., Lanis, J.M., Alexeev, E.E., Wang, R.X., Onyiah, J.C., Kominsky, D.J., and Colgan, S.P. (2017). Microbial-Derived Butyrate Promotes Epithelial Barrier Function through IL-10 Receptor–Dependent Repression of Claudin-2. The Journal of Immunology 199, 2976–2984. 10.4049/jimmunol.1700105.

48. Dang, H., Feng, P., Zhang, S., Peng, L., Xing, S., Li, Y., Wen, X., Zhou, L., Goswami, S., Xiao, M., et al. (2025). Maternal gut microbiota influence stem cell function in offspring. Cell Stem Cell 32, 246–262 e248. 10.1016/j.stem.2024.10.003.

49. Schlosser, S., Ullrich, A.L., Modares, N.F., Schmitz, M.A., Schoneich, J., Zhang, K., Richter, I., Robrahn, L., Schraven, S., Nagai, J.S., et al. (2025). Salmonella infection accelerates postnatal maturation of the intestinal epithelium. Proc Natl Acad Sci U S A 122, e2403344122. 10.1073/pnas.2403344122.

50. Kim, D., Langmead, B., and Salzberg, S.L. (2015). HISAT: a fast spliced aligner with low memory requirements. Nature Methods 12, 357–360. 10.1038/nmeth.3317.

51. Liao, Y., Smyth, G.K., and Shi, W. (2014). featureCounts: an efficient general purpose program for assigning sequence reads to genomic features. Bioinformatics 30, 923–930. 10.1093/bioinformatics/btt656.

52. Love, M.I., Huber, W., and Anders, S. (2014). Moderated estimation of fold change and dispersion for RNA-seq data with DESeq2. Genome Biology 15, 550. 10.1186/s13059-014-0550-8.

53. Shannon, P., Markiel, A., Ozier, O., Baliga, N.S., Wang, J.T., Ramage, D., Amin, N., Schwikowski, B., and Ideker, T. (2003). Cytoscape: A Software Environment for Integrated Models of Biomolecular Interaction Networks. Genome Research 13, 2498–2504.

54. Cline, M.S., Smoot, M., Cerami, E., Kuchinsky, A., Landys, N., Workman, C., Christmas, R., Avila-Campilo, I., Creech, M., Gross, B., et al. (2007). Integration of biological networks and gene expression data using Cytoscape. Nat Protoc 2, 2366–2382. 10.1038/nprot.2007.324.

55. Doncheva, N.T., Morris, J.H., Gorodkin, J., and Jensen, L.J. (2019). Cytoscape StringApp: Network Analysis and Visualization of Proteomics Data. J Proteome Res 18, 623–632. 10.1021/acs.jproteome.8b00702.

56. Enright, A.J., Van Dongen, S., and Ouzounis, C.A. (2002). An efficient algorithm for large-scale detection of protein families. Nucleic Acids Research 30, 1575–1584. 10.1093/nar/30.7.1575.

57. Balwierz, P.J., Pachkov, M., Arnold, P., Gruber, A.J., Zavolan, M., and Van Nimwegen, E. (2014). ISMARA: automated modeling of genomic signals as a democracy of regulatory motifs. Genome Research 24, 869–884. 10.1101/gr.169508.113.

58. Newman, A.M., Steen, C.B., Liu, C.L., Gentles, A.J., Chaudhuri, A.A., Scherer, F., Khodadoust, M.S., Esfahani, M.S., Luca, B.A., Steiner, D., et al. (2019). Determining cell type abundance and expression from bulk tissues with digital cytometry. Nature Biotechnology 37, 773–782. 10.1038/s41587-019-0114-2.

59. Haber, A.L., Biton, M., Rogel, N., Herbst, R.H., Shekhar, K., Smillie, C., Burgin, G., Delorey, T.M., Howitt, M.R., Katz, Y., et al. (2017). A single-cell survey of the small intestinal epithelium. Nature 551, 333–339. 10.1038/nature24489.

60. Ewels, P.A., Peltzer, A., Fillinger, S., Alneberg, J., Patel, H., Wilm, A., Garcia, M.U., Di Tommaso, P., and Nahnsen, S. (2019). nf-core: Community curated bioinformatics pipelines. bioRxiv, 610741. 10.1101/610741.

61. Krueger, F., and Andrews, S.R. (2011). Bismark: a flexible aligner and methylation caller for Bisulfite-Seq applications. Bioinformatics 27, 1571–1572. 10.1093/bioinformatics/btr167.

62. Akalin, A., Kormaksson, M., Li, S., Garrett-Bakelman, F.E., Figueroa, M.E., Melnick, A., and Mason, C.E. (2012). methylKit: a comprehensive R package for the analysis of genome-wide DNA methylation profiles. Genome Biology 13, R87. 10.1186/gb-2012-13-10-r87.

63. Jühling, F., Kretzmer, H., Bernhart, S.H., Otto, C., Stadler, P.F., and Hoffmann, S. (2016). metilene: fast and sensitive calling of differentially methylated regions from bisulfite sequencing data. Genome Research 26, 256–262.

64. Korthauer, K., Chakraborty, S., Benjamini, Y., and Irizarry, R.A. (2019). Detection and accurate false discovery rate control of differentially methylated regions from whole genome bisulfite sequencing. Biostatistics 20, 367–383. 10.1093/biostatistics/kxy007.

65. Duttke, S.H., Chang, M.W., Heinz, S., and Benner, C. (2019). Identification and dynamic quantification of regulatory elements using total RNA. Genome Research 29, 1836–1846.

66. Ewels, P.A., Peltzer, A., Fillinger, S., Patel, H., Alneberg, J., Wilm, A., Garcia, M.U., Di Tommaso, P., and Nahnsen, S. (2020). The nf-core framework for community-curated bioinformatics pipelines. Nat Biotechnol 38, 276–278. 10.1038/s41587-020-0439-x.

67. Love, M.I., Huber, W., and Anders, S. (2014). Moderated estimation of fold change and dispersion for RNA-seq data with DESeq2. Genome Biol 15, 550. 10.1186/s13059-014-0550-8.

68. Yu, G., Wang, L.G., and He, Q.Y. (2015). ChIPseeker: an R/Bioconductor package for ChIP peak annotation, comparison and visualization. Bioinformatics 31, 2382–2383. 10.1093/bioinformatics/btv145.

69. Wu, T., Hu, E., Xu, S., Chen, M., Guo, P., Dai, Z., Feng, T., Zhou, L., Tang, W., Zhan, L., et al. (2021). clusterProfiler 4.0: A universal enrichment tool for interpreting omics data. Innovation (Camb) 2, 100141. 10.1016/j.xinn.2021.100141.

70. Fuhrer, T., Heer, D., Begemann, B., and Zamboni, N. (2011). High-throughput, accurate mass metabolome profiling of cellular extracts by flow injection-time-of-flight mass spectrometry. Anal Chem 83, 7074–7080. 10.1021/ac201267k.

71. Djoumbou Feunang, Y., Eisner, R., Knox, C., Chepelev, L., Hastings, J., Owen, G., Fahy, E., Steinbeck, C., Subramanian, S., Bolton, E., et al. (2016). ClassyFire: automated chemical classification with a comprehensive, computable taxonomy. Journal of Cheminformatics 8, 61. 10.1186/s13321-016-0174-y.

72. Grün, D., Lyubimova, A., Kester, L., Wiebrands, K., Basak, O., Sasaki, N., Clevers, H., and van Oudenaarden, A. (2015). Single-cell messenger RNA sequencing reveals rare intestinal cell types. Nature 525, 251–255. 10.1038/nature14966.

73. Moor, A.E., Harnik, Y., Ben-Moshe, S., Massasa, E.E., Rozenberg, M., Eilam, R., Bahar Halpern, K., and Itzkovitz, S. (2018). Spatial Reconstruction of Single Enterocytes Uncovers Broad Zonation along the Intestinal Villus Axis. Cell 175, 1156–1167.e1115. 10.1016/j.cell.2018.08.063.

